# Integrative Multiomics to Dissect the Lung Transcriptional Landscape of Pulmonary Arterial Hypertension

**DOI:** 10.1101/2023.01.12.523812

**Authors:** Jason Hong, Brenda Wong, Christopher J. Rhodes, Zeyneb Kurt, Tae-Hwi Schwantes-An, Elizabeth A. Mickler, Stefan Gräf, Mélanie Eyries, Katie A. Lutz, Michael W. Pauciulo, Richard C. Trembath, David Montani, Nicholas W. Morrell, Martin R. Wilkins, William C. Nichols, David-Alexandre Trégouët, Micheala A. Aldred, Ankit A. Desai, Rubin M. Tuder, Mark W. Geraci, Mansoureh Eghbali, Robert S. Stearman, Xia Yang

**Author notes:** **Correspondence** Jason Hong, M.D., Ph.D., Division of Pulmonary and Critical Care Medicine, David Geffen School of Medicine at UCLA, 200 UCLA Medical Plaza, Suite 365-B, Box 951693, Los Angeles, CA 90095. Co-senior authors (Mansoureh Eghbali, Robert S. Stearman, Xia Yang).

## Abstract

Pulmonary arterial hypertension (PAH) remains an incurable and often fatal disease despite currently available therapies. Multiomics systems biology analysis can shed new light on PAH pathobiology and inform translational research efforts. Using RNA sequencing on the largest PAH lung biobank to date (96 disease and 52 control), we aim to identify gene co-expression network modules associated with PAH and potential therapeutic targets. Co-expression network analysis was performed to identify modules of co-expressed genes which were then assessed for and prioritized by importance in PAH, regulatory role, and therapeutic potential via integration with clinicopathologic data, human genome-wide association studies (GWAS) of PAH, lung Bayesian regulatory networks, single-cell RNA-sequencing data, and pharmacotranscriptomic profiles. We identified a co-expression module of 266 genes, called the pink module, which may be a response to the underlying disease process to counteract disease progression in PAH. This module was associated not only with PAH severity such as increased PVR and intimal thickness, but also with compensated PAH such as lower number of hospitalizations, WHO functional class and NT-proBNP. GWAS integration demonstrated the pink module is enriched for PAH-associated genetic variation in multiple cohorts. Regulatory network analysis revealed that BMPR2 regulates the main target of FDA-approved riociguat, GUCY1A2, in the pink module. Analysis of pathway enrichment and pink hub genes (i.e. ANTXR1 and SFRP4) suggests the pink module inhibits Wnt signaling and epithelial-mesenchymal transition. Cell type deconvolution showed the pink module correlates with higher vascular cell fractions (i.e. myofibroblasts). A pharmacotranscriptomic screen discovered ubiquitin-specific peptidases (USPs) as potential therapeutic targets to mimic the pink module signature. Our multiomics integrative study uncovered a novel gene subnetwork associated with clinicopathologic severity, genetic risk, specific vascular cell types, and new therapeutic targets in PAH. Future studies are warranted to investigate the role and therapeutic potential of the pink module and targeting USPs in PAH.

## Introduction

Pulmonary arterial hypertension (PAH) remains an incurable and often fatal disease characterized by irreversible vascular remodeling. Despite the identification of many candidate drugs in the preclinical stage, effective therapies that reverse the underlying disease process are still lacking. A deeper understanding of the molecular and cellular mechanisms in PAH lung tissue is needed to bridge this translational gap.

Data-driven transcriptome-wide studies of PAH lungs have uncovered genes and pathways differentially expressed in PAH(1, 2). However, whether such findings are robust, causal, and cell-specific in disease pathogenesis remain unknown since lung samples are usually from limited numbers of advanced stage PAH patients at the bulk tissue level. Furthermore, typical gene-level differential expression analysis may not reveal upstream causal and regulatory genes and pathways(3, 4). A rigorous systems-level examination of altered transcriptomes in PAH lungs integrating different omics data types is needed to advance our understanding of PAH pathobiology and help inform potential causal genes, regulatory networks and pathways, and therapeutic targets to facilitate translational efforts.

In this study which leverages the transcriptional landscape of a large biorepository of PAH lungs (96 disease vs 52 control), we dissect the gene networks of PAH lungs using an integrative multiomic and systems biology approach to uncover a module of co-expressed genes associated with clinicopathologic severity, genetic risk, and vascular cell specificity in PAH, and further identify novel therapeutic targets by a pharmacotranscriptomic screen for future preclinical studies.

## Methods

### RNA sequencing and differential expression analysis

Patient enrollment and the standardized tissue-processing protocol for PHBI have been previously described(1, 5). Paired-end 75 base-pair RNA sequencing (RNA-seq) was performed on all available PHBI lung samples using an Illumina sequencer. Samples were sequenced in two batches. Sequencing depth was 20-25 million reads per sample in one batch and 15-20 million reads per sample in the other batch. Reads were mapped to the UCSC human reference genome (version hg19) using STAR(6). Transcripts were assembled and quantified using StringTie(7). Transcript-level abundance estimates were imported and summarized into a counts matrix using tximport(8) which was then input into DESeq2 (9) for differential gene expression analysis using a negative binomial generalized linear model. Potential outliers and batch effects of different covariates (i.e. sequencing batch, sex, age, ethnicity) were assessed by hierarchical clustering and principal component analysis. Two patients with WHO Group IV pulmonary hypertension were excluded from this analysis. Sequencing batch was adjusted for in the DESeq2 model. Sex-stratified differential expression analysis was also performed. Differentially expressed genes with FDR < 0.05 were considered statistically significant.

### Weighted Gene Co-Expression Network Analysis (WGCNA)

WGCNA v. 1.69 R package was used to identify modules of co-expressed genes in PHBI lung RNA-seq samples. We first performed variance stabilizing transformation of the counts matrix using DESeq2. We then adjusted for the effect of the two sequencing batches using an empirical Bayes framework through the ComBat function of the sva v. 3.34 R package. The batch-corrected expression matrix was then evaluated for potential outlier samples by hierarchal clustering. Two samples were identified as outliers and were removed from downstream analyses: a 55 year-old female control and a 14 year-old male patient with idiopathic PAH. The bottom 25% of genes with the least variation across samples were filtered out. Known PAH genes that were filtered out at this step were added back to the expression matrix (48 out of 582 PAH genes retrieved from disease-gene databases DisGeNET (10) and Comparative Toxicogenomics Database (11) using the Harmonizome portal (12)). A total of 17,564 genes were then included in downstream WGCNA steps. A soft-thresholding power of 3 was selected to power the correlation of genes with the assumption that raising the correlation to a specific power will reduce the noise of the correlations in the adjacency matrix. A soft-thresholding power of 3 was selected to optimize both the scale-free topology index (R^2^ > 0.8) and mean connectivity (k = 205). To minimize effects of noise and spurious associations, the adjacency matrix was transformed into a Topological Overlap Matrix (TOM), and the corresponding dissimilarity matrix was calculated. Hierarchical clustering was then performed on the dissimilarity matrix after which genes were split into 25 modules using the cutreeDynamic function of the dynamicTreeCut R package using the following parameters: DeepSplit = 2, pamRespectsDendro = FALSE, cutHeight = 0.99, minClusterSize = 30. Module eigengenes representing the first principal component of a given module in a given single dataset (i.e. PHBI lung dataset) were calculated using the moduleEigengenes function of WGCNA. Given that dynamicTreeCut may identify modules whose expression profiles are similar in which their genes are highly co-expressed, such modules were assessed for and merged using the following step per the WGCNA analysis pipeline: hierarchical clustering was performed on the dissimilarity of module eigengenes after which similar modules were merged using the mergeCloseModules function of WGCNA using a cutHeight of 0.15. Merging of similar modules yielded 20 final modules which were the used for downstream analyses. The strongest pairwise gene-gene connection (ANTXR1 and SFRP4) within the pink module was identified by comparing TOM values across all 33,732 pairs of pink genes.

### Pathway enrichment analysis

Gene set enrichment analysis (GSEA) using R package fgsea v1.18.0 was performed to identify enriched pathways in the PAH lung signature. The PAH signature represents the differential transcriptome between PAH and control. Genes were ordered by the Wald statistic as determined by DESeq2 of PAH vs. control. Per the GSEA algorithm, genes were not filtered (i.e. by expression, variance, or by a statistical threshold for differential expression) prior to GSEA. The PAH signature was tested for enrichment in Hallmark pathways from Molecular Signature Database (13), as well as in co-expression modules. Sex-stratified analysis was performed using the sex-stratified differential expression analysis results. GSEA was also performed on select co-expression module signatures (i.e. pink, royalblue, greenyellow). Module signatures were defined as the correlation of the module eigengene with the expression of genes across the transcriptome. Similar to the PAH signature analysis, genes were not filtered prior to GSEA. Module signatures were tested for enrichment in Biological Processes from Gene Ontology (GO) (14), Hallmark pathways from Molecular Signature Database (13), and/or known cell-type signatures from Azimuth(15). Enrichment in pathways with FDR < 0.05 were considered statistically significant.

### Module-trait correlation analysis

Pearson correlations of module eigengenes with clinical and pathologic characteristics were computed in order to prioritize modules by importance in PAH. Dichotomous categorical variables were coded 0 and 1. WHO functional class was obtained from the New York Heart Association (NYHA) class recorded immediately pre-transplant (i.e. day of transplant) and was coded as integers 1, 2, 3 and 4. If the immediate pre-transplant NYHA class was unavailable for a given patient, then the WHO functional class from their most recent clinic visit was used. Number of hospitalizations due to PAH were counted between the time of diagnostic RHC and lung transplantation. Presence of right heart failure signs such as ascites or leg swelling were recorded at the time of enrollment in the study (i.e. just prior to lung transplantation). Lab values (i.e. NT-proBNP and creatinine) were obtained from the most recent blood draw prior to transplant. The most recent pulmonary function testing and right heart catheterization results prior to transplant were used. Intima and intima plus media thickness were measured by a lung pathologist on explant histological tissue sections by morphometric analysis of volume density of pulmonary arteries. REVEAL lite scores were calculated as per Benza et al(16) using values of NT-proBNP or BNP, six-minute walk distance, WHO functional class, systolic blood pressure, heart rate, and creatinine. Values were obtained from the most recent available assessment prior to transplant. A score of zero was assigned for missing individual assessments as per Benza et al(16). *P* values < 0.05 were considered statistically significant. To minimize type II error and potential false negative results, results were interpreted using nominal *P* values given that our module-trait correlation analysis was intended to be exploratory and hypothesis-generating rather than to confirm an a prior hypothesis about module-trait correlations. Results will need to be confirmed in future targeted studies. Multiple testing correction (n=260 comparisons from 13 traits and 20 modules) was also performed and correlations with FDR < 0.05 are shown in a supplemental figure.

### Genome-wide association study (GWAS) enrichment analysis

Enrichment of modules for PAH GWAS single-nucleotide polymorphisms (SNPs) were assessed using two distinct computational methods, MAGMA(17) and GSA-SNP2(18), across four independent PAH GWAS cohorts totaling 11,744 individuals(19, 20): the US PAH Biobank (PAHB), French Pulmonary Hypertension Allele-Associated Risk (PHAAR), British Heart Foundation Pulmonary Arterial Hypertension (BHFPAH), and UK National Institute for Health Research BioResource (NIHRBR) cohorts. SNPs were mapped to genes by chromosomal proximity (within 20 kilobases from the 5’ or 3’ ends of a gene) and genes were scored for association with PAH based on disease-SNP *P*-value associations from GWAS summary statistics. SNPs were not filtered (i.e. by a specified statistical threshold) prior to input into MAGMA and GSA-SNP2. Gene scores were then used in competitive gene-set analyses to identify module enrichment for PAH common genetic variation. To aggregate genetic variants into a gene score, the mean χ^2^ statistic and the log-minimum GWAS *P* value for all SNPs localizing to a gene were used as per MAGMA(17) and GSA-SNP2(18), respectively. To determine significance, MAGMA uses a linear mixed model whereas GSA-SNP2 uses a standard normal distribution. Both methods adjust for gene size and gene density (the number of SNPs assigned to a given gene). The default statistical results were reported for MAGMA (*P* value) and GSA-SNP2 (false discovery rate). MAGMA *P* values were not corrected post-hoc for multiple testing given that this analysis was intended to be exploratory and hypothesis-generating rather than to confirm an a prior hypothesis about GWAS enrichment. Results will need to be confirmed in future targeted studies.

### Bayesian gene regulatory network analysis

In a complementary approach to co-expression analysis to infer co-regulation, we employed Bayesian network (BN) analysis to build a gene regulatory network. Specifically, BNs were constructed using Reconstructing Integrative Molecular Bayesian Network (RIMBANet)(21). For this method, 1000 networks were generated from different random seed genes using continuous and discrete expression data derived from transcriptomes from either GSE23546 (n = 1343) (22), PHBI (n = 146), or GTEx v8 (n = 577) (23). Whole lung-specific cis eQTLs from GTEx v8 (23) and transcription factor-target gene data from HTRI (24), TRRUST (25), and PAZAR (26) databases were used as priors. Then, the final network for each of the 3 datasets was obtained by taking a consensus network from the 1000 randomly generated networks whereby only edges that passed a probability of >30% across the 1000 BNs were kept. Finally, the union of the 3 networks was taken to create a combined gene regulatory network derived from a total of 2,066 human lungs. The network was visualized in Cytoscape(27) where nodes represent genes and edges represent inferred directional gene-gene regulation. Node positions were determined by a prefuse force-directed algorithm. Genes from co-expression modules were projected onto this regulatory network with different color nodes representing module membership. Known PAH genes from disease-gene databases (Comparative Toxicogenomics Database(11) and DisGeNET(10)) were also projected onto the network. Local hub genes of a particular subnetwork (i.e. pink subnetwork) whose neighboring nodes are enriched for genes in the gene set of interest (i.e. pink gene set) were determined by Key Driver Analysis in the Mergeomics R package(4, 28, 29).

### Analysis of public transcriptomic datasets

Public transcriptomic datasets were queried for expression of selected genes (i.e. *GUCY1A2*, *ANTXR1, USP28,* and *USP12*). *GUCY1A2* expression was obtained from two independent RNA-seq datasets: CRISPR/Cas9-induced monoallelic mutations in BMPR2 (n = 6) vs wild-type control (n = 3) human umbilical vein endothelial cells (HUVEC)(30) and endothelial cells derived from induced pluripotent stem cells (iPSCs) of patients with hereditary PAH (HPAH) due to BMPR2 mutations (n = 5) vs control (n = 3)(31). Sequence Read Archive (SRA) data was downloaded from the NIH Gene Expression Omnibus (GEO) database. SRA files were converted to FASTQ files using the SRA Toolkit. Sequences were aligned to the GENCODE human reference genome (v. 32) using HISAT2(7) and transcripts were assembled and quantified using StringTie(7). DESeq2(9) was used to perform differential expression analysis and determine *P* values. *ANTXR1* expression in fibroblasts was obtained from an scRNAseq dataset comparing 3 PAH vs 6 control lungs (32). *ANTXR1* expression counts were averaged across cells within the fibroblast cluster for each sample. Note, myofibroblasts were not subclustered in this dataset. Wilcoxon rank-sum test was used to determine differential expression of *ANTXR1* in PAH vs. control samples. The expression of *USP28* and *USP12* was obtained from a human whole lung microarray of 15 PAH patients vs 11 controls(33) deposited in the NIH GEO database. *P* values were obtained from NCBI’s GEO2R.

### Quantitative polymerase chain reaction (qPCR)

Pulmonary arterial adventitial cells isolated from idiopathic PAH and control lungs were obtained from PHBI and grown in Human Vascular Smooth Muscle Cell Basal Medium (M231500, ThermoFisher) supplemented with Smooth Muscle Growth Supplement (S00725, ThermoFisher) and Antibiotic-Antimycotic 100X (15240096, ThermoFisher). Both PAH and control cells originated from the lungs of white women age 29 and 33, respectively. Cells were collected in TRIzol (15596018, ThermoFisher) once grown to 100% confluency between passages 6 to 10. Different passages served as biological replicates. RNA was extracted from cells through a series of washes using chloroform (364320025, ThermoFisher), isopropanol (I9516, MillaporeSigma), and 70% ethanol (459844, MillaporeSigma). RNA was resuspended in DEPC-Treated Water (AM9922,ThermoFisher) and then converted to cDNA using a High-Capacity cDNA Reverse Transcription Kit (4368814, ThermoFisher) and a Bio-Rad S1000 Thermal Cycler. A qCPR was run using cDNA, DEPC water, PowerUp™ SYBR™ Green Master Mix (A25743, ThermoFisher) and primers on a Bio-Rad CFX Connect Real-Time PCR Detection System. Primers were used for ANTXR1 (Forward: GAGGAAACGGCTTCCGACAT, Reverse: GAGTGCAGCTTTCATGCCAA) and housekeeping gene RPLP0 (Forward: CAGGTGTTCGACAATGGCAG, Reverse: ACAAGGCCAGGACTCGTTTG).

### Deconvolution

To serve as a cell type reference for deconvolution, we integrated seven publicly available human lung single-cell RNAseq datasets(34–40) and identified 37 cell-type clusters using known marker genes from the literature. Within each cell-type cluster, the average expression of gene counts was calculated across cells for each individual sample to create a cell-type signature for each of the seven datasets. PHBI bulk transcriptomes were deconvoluted with CIBERSORTx(41) with cell-type signatures from each of the seven datasets as a reference. The resulting cell fractions using each of the seven dataset-specific reference signatures served as technical replicates. These technical replicates were then averaged to determine the final estimated cell fractions for each lung sample. Pearson correlation of deconvoluted cell type fractions with PAH vs control status (coded 1 and 0, respectively) was calculated across PHBI lung samples. Wilcoxon rank-sum test was performed on cell fractions between PAH vs. control samples for vascular cell types. Similar to module-trait correlation analysis, modules were correlated with cell fractions by calculating Pearson correlations of module eigengenes with deconvoluted cell fractions across samples.

### Pharmacotranscriptomic analysis

Genes differentially expressed between PAH and control in select co-expression modules (i.e. pink, royalblue, greenyellow) were queried against the full Connectivity Map(42) (CMap) database of perturbagen expression signatures induced in human cell lines(42). A less stringent statistical threshold for differentially expressed genes (*P* value < 0.05) were used as previously described(43) to ensure an adequate number of query signature genes to perform the CMap analysis. A total of 8,559 pharmacologic and genetic perturbagens were screened including both gene overexpression and knockdown by short hairpin RNA (shRNA). Pattern-matching algorithms assessed each reference perturbagen profile for the direction and strength of connectivity with the query signature by a score range of -100 to +100. The summary score was used across 9 cell lines. Perturbagens with strongly positive connectivity scores indicate highly similar signatures that mimic that of the query whereas those perturbagens with strongly negative scores indicate signatures that strongly reverse that of the query (i.e. genes that are differentially upregulated in the module query are decreased by the perturbagen or vice versa). We also assessed the connectivity scores of a total of 171 CMap classes defined as groups of pharmacologic or genetic CMap perturbagens that share the same mechanism of action or biological function.

The pink signature of differentially expressed genes (PAH vs. control; *P* value < 0.05) was also queried against the CRISPR knockout (KO) consensus signature database of Library of Integrated Network-based Cellular Signatures (LINCS) L1000 using SigCom LINCS(44). A total of 7,502 genes were screened by CRISPR KO from this database. SigCom outputs a Z score which indicates the degree to which the CRISPR KO signature mimics or reverses the query signature (i.e. pink) by highly positive or negative scores, respectively. The expression of 12,327 genes were profiled and ranked for each CRISPR KO gene where lowly and highly ranked genes indicate downregulation and upregulation, respectively, in the CRISPR KO vs. control.

## Results

### Characterization of PHBI cohort

RNA sequencing was performed on a total of 96 explant lungs with pulmonary hypertension (PH) collected at the time of lung transplantation and 52 control lungs from the Pulmonary Hypertension Breakthrough Initiative (PHBI) (Table 1 and Figure 1A). WHO group 1 PAH patients consisted of 94 of 96 PH subjects of which the most common subtypes were idiopathic PAH (IPAH) and associated PAH (APAH) (43% and 40%, respectively). The majority of PH patients were WHO functional class III or IV, had significantly impaired hemodynamics by right heart catheterization with a mean PVR of 12.7±7.4, and were receiving triple PAH-targeted therapy (73%) including prostacyclin infusion therapy (85%). Unsupervised hierarchical clustering and principal component analysis (PCA) showed separation between PH and control samples suggesting overall differences in the transcriptome between the two groups (Figures 1A-1B). Neither approach showed distinct separation of samples by PAH subgroup suggesting relative lack of subgroup-specific transcriptional heterogeneity. Furthermore, we did not observe significant clustering by age, sex, or race among PH or control samples. Moreover, samples did not cluster by transplant center of tissue origin nor treatment group (Supplemental Figure 1). Given the female predominance of PH subjects compared to control, sex-stratified analyses were performed where appropriate. Two outliers (1 IPAH and 1 control) were removed from downstream analyses (Figures 1A-1B) since network analysis and module detection can be biased by outlier samples(45).

**Table 1:**
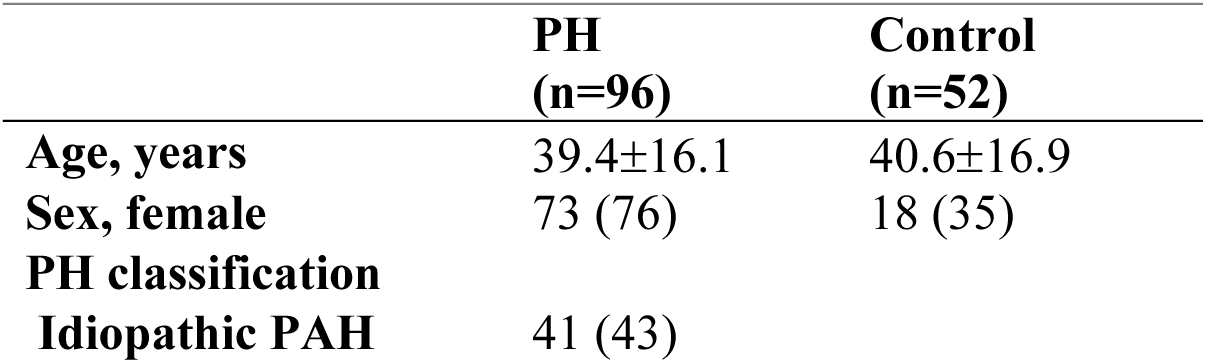
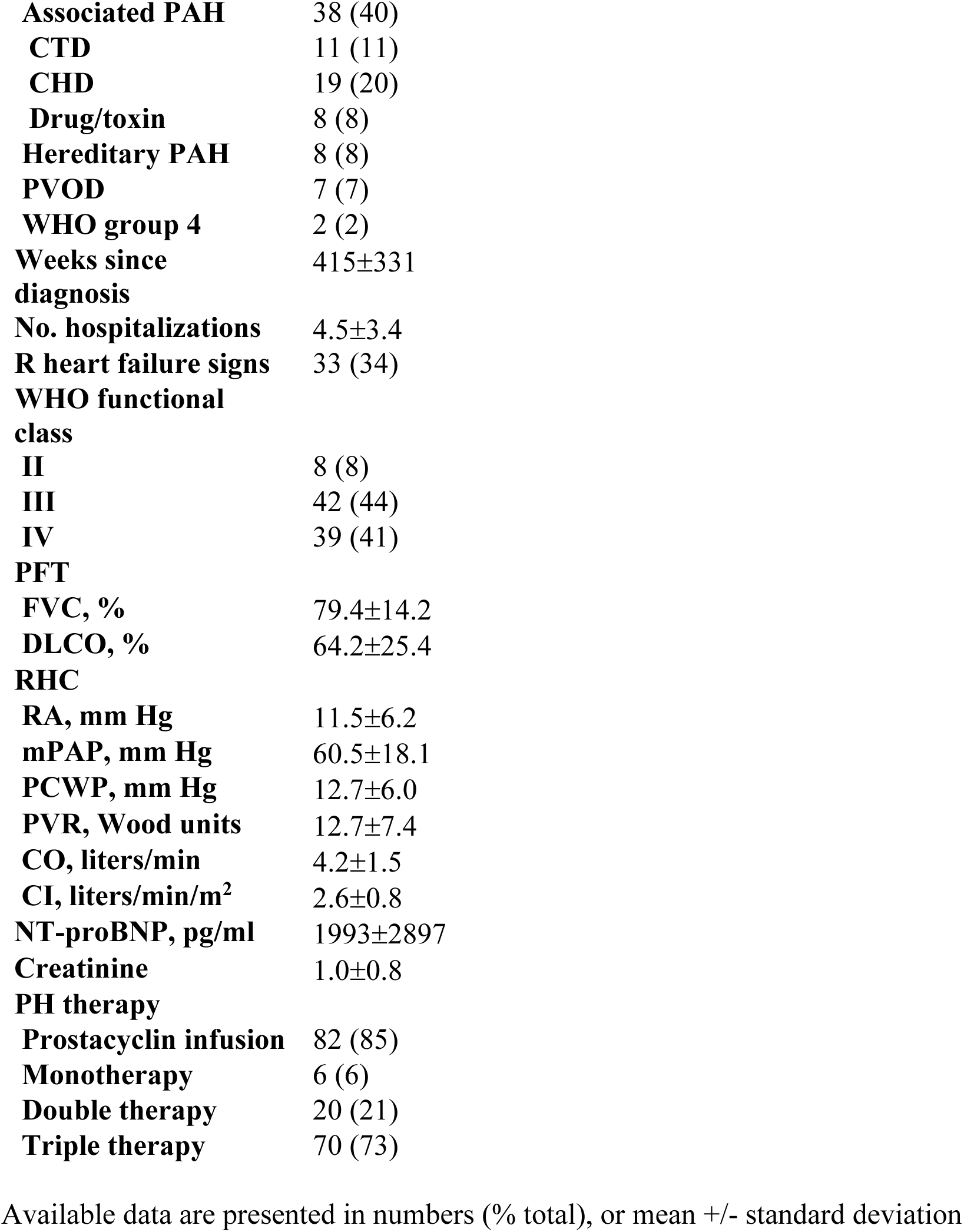
Patient characteristics.

|  | **PH**<br>**(n=96)** | **Control**<br>**(n=52)** |
| --- | --- | --- |
| **Age, years** | 39.4±16.1 | 40.6±16.9 |
| **Sex, female** | 73 (76) | 18 (35) |
| **PH classification** |  |  |
| **Idiopathic PAH** | 41 (43) |  |

|  |  |
| --- | --- |
| **Associated PAH** | 38 (40) |
| **CTD** | 11 (11) |
| **CHD** | 19 (20) |
| **Drug/toxin** | 8 (8) |
| **Hereditary PAH** | 8 (8) |
| **PVOD** | 7 (7) |
| **WHO group 4** | 2 (2) |
| **Weeks since diagnosis** | 415±331 |
| **No. hospitalizations** | 4.5±3.4 |
| **R heart failure signs** | 33 (34) |
| **WHO functional class** |  |
| **II** | 8 (8) |
| **III** | 42 (44) |
| **IV** | 39 (41) |
| **PFT** |  |
| **FVC, %** | 79.4±14.2 |
| **DLCO, %** | 64.2±25.4 |
| **RHC** |  |
| **RA, mm Hg** | 11.5±6.2 |
| **mPAP, mm Hg** | 60.5±18.1 |
| **PCWP, mm Hg** | 12.7±6.0 |
| **PVR, Wood units** | 12.7±7.4 |
| **CO, liters/min** | 4.2±1.5 |
| **CI, liters/min/m<sup>2</sup>** | 2.6±0.8 |
| **NT-proBNP, pg/ml** | 1993±2897 |
| **Creatinine** | 1.0±0.8 |
| **PH therapy** |  |
| **Prostacyclin infusion** | 82 (85) |
| **Monotherapy** | 6 (6) |
| **Double therapy** | 20 (21) |
| **Triple therapy** | 70 (73) |

Available data are presented in numbers (% total), or mean +/- standard deviation

**Figure 1:**
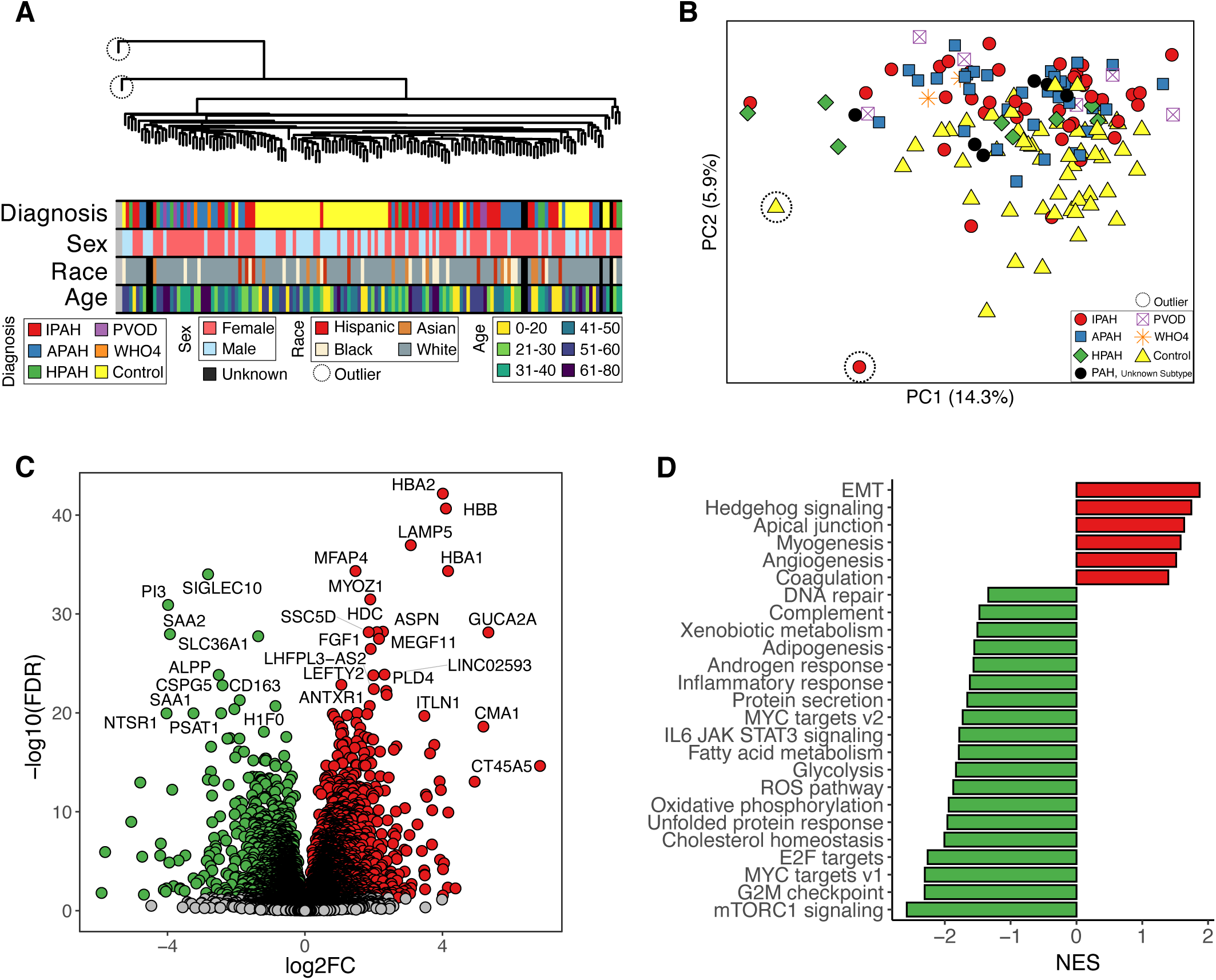
The lung transcriptome is significantly altered in PAH. (A) Unsupervised hierarchical clustering of PHBI transcriptomes: 17,567 genes after filtering the bottom 25% of genes with the least variation across samples. Samples are annotated by age, sex, race, and diagnosis. Two outliers are circled. (B) PCA plot showing PCA of all 23,355 detected genes where samples are colored by diagnosis. The same two outliers as in (A) are circled. (C) Volcano plot showing upregulated genes colored in red and downregulated genes colored in green. Grey dots indicate genes with FDR ≥ 0.05. (D) Bar plot showing GSEA results using the Hallmark pathway database. Pathways enriched in genes upregulated in PAH are colored in red and pathways enriched in downregulated genes in green. Only pathways with FDR < 0.05 are shown. IPAH = idiopathic PAH; APAH = associated PAH; HPAH = hereditary PAH; PVOD = pulmonary veno-occlusive disease; WHO4 = WHO group 4 PAH; FDR = false discovery rate; log2FC = log2 fold change; NES = normalized enrichment score; EMT = epithelial- mesenchymal transition.

### The lung transcriptome is significantly altered in PAH

Consistent with the separation observed between PH and control samples by hierarchical clustering and PCA, differential expression analysis between PAH and control samples yielded 5,253 differentially expressed genes (DEGs; FDR < 0.05) consisting of 22% of the transcriptome of which 2,719 were upregulated and 2,534 were downregulated (Figure 1C). The top upregulated genes were *HBA2*, *HBB*, *LAMP5*, *HBA1*, and *MFAP4* whereas the top downregulated genes were *SIGLEC10*, *PI3*, *SAA2*, *SLC36A1*, and *ALPP*. Epithelial-mesenchymal transition (EMT) was the top enriched pathway among genes upregulated in PAH whereas mTORC1 signaling was the top pathway among downregulated genes (Figure 1D).

### Co-expression network analysis reveals modules associated with PAH severity

We then used weighted gene co-expression network analysis (WGCNA)(46) across all samples to dissect the lung transcriptome into clusters based on gene co-expression, referred to as modules. Modules organize transcriptional changes of individual genes into clusters which represent co-regulation or shared biological functions(47) (Figure 2A-B). We identified 20 gene co-expression modules with a median size of 141 genes (Figure 2C). The expression of genes within a module can be summarized by their eigengene which represents the first principal component of gene expression in the module. Correlation of clinicopathologic characteristics with module eigengenes revealed the pink module of 266 genes to have the most notable pattern of associations (Figure 2D). The pink module was not only strongly associated with PAH diagnosis (Figures 2D-2E, Supplemental Figure 2) but also with physiologic, hemodynamic, and pathologic markers of disease severity based on pulmonary function testing, right heart catheterization, and histologic analysis of vascular remodeling by morphometry. Specifically, the pink module correlated with reduced diffusing capacity for carbon monoxide (DLCO), elevated mean pulmonary artery pressure (mPAP), elevated pulmonary vascular resistance (PVR), and increased intima or intima plus media thickness of pulmonary arteries. However, the pink module also correlated with clinical characteristics and blood tests suggestive of compensated disease, such as lower number of hospitalizations due to PAH, signs of right heart failure, WHO functional class, NT-proBNP, creatinine, and REVEAL lite score. Among other modules positively correlated with PAH diagnosis, the royalblue module of 98 genes shared a similar pattern of clinicopathologic correlation as with the pink module. The greenyellow module of 290 genes had the highest negative correlation with PAH diagnosis (Figure 2F). Pink and royalblue were also the top two modules most strongly enriched for genes upregulated in PAH lungs and greenyellow was most strongly enriched for downregulated genes (Figure 2G, Supplemental Figure 3).

**Figure 2:**
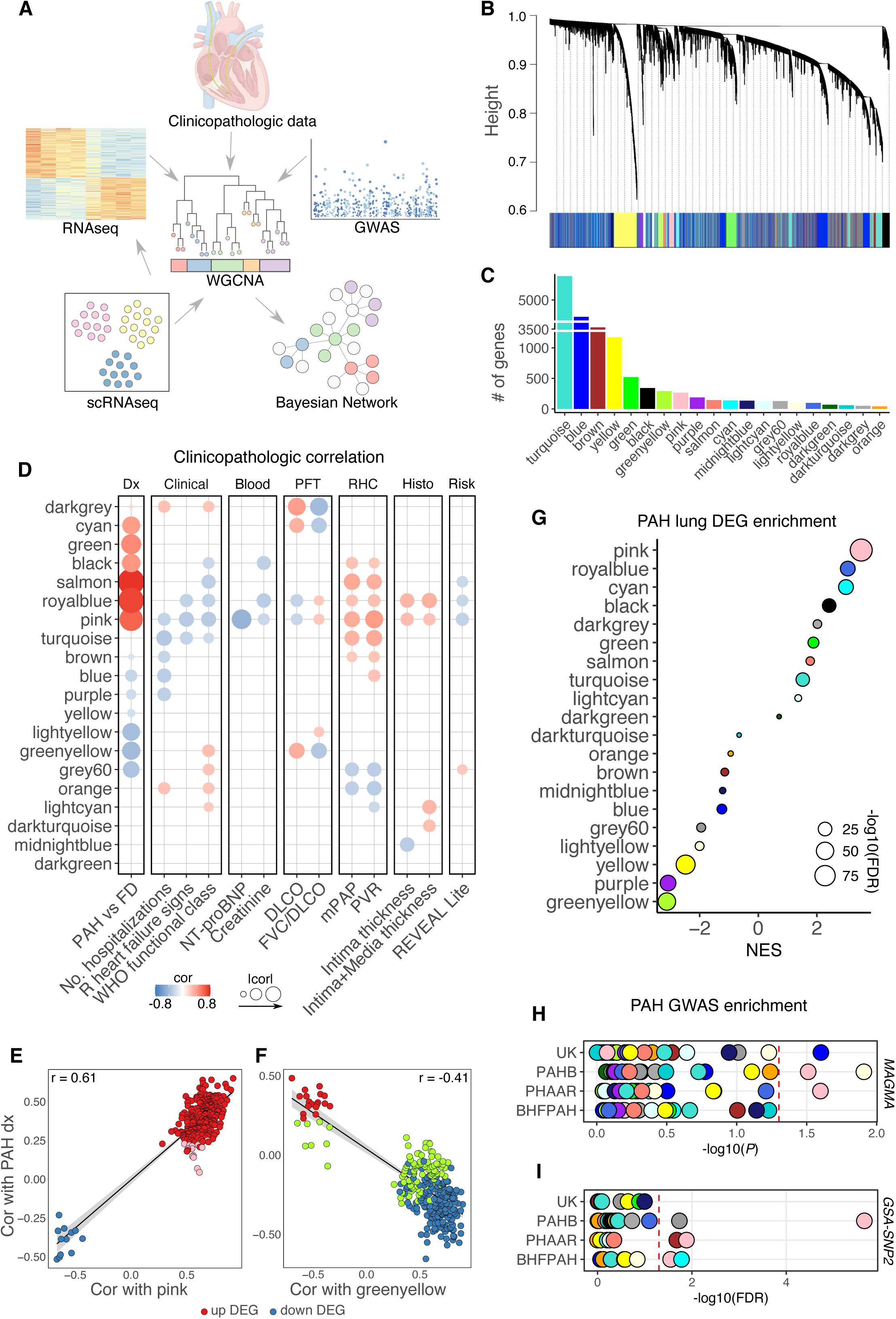
Co-expression network analysis reveals modules associated with PAH severity and genetic variants. (A) Schematic of integrative analytical strategy centered around co-expression modules. (B) Gene clustering dendrogram as determined by WGCNA with color module assignments shown at the bottom. (C) Bar plot showing number of genes in each module. (D) Heatmap showing significant (*P* < 0.05) Pearson correlations of module eigengenes with clinical and pathologic characteristics where red and blue indicate positive and negative correlation, respectively. Larger size dots indicate stronger correlation. No. hospitalizations indicates number of hospitalizations due to PAH between the time of diagnostic RHC and lung transplantation. R heart failure signs indicate signs of right heart failure such as ascites or leg swelling. Intima and intima plus media thickness were determined by morphometric analysis of volume density of pulmonary arteries in histological lung sections. (E-F) Scatter plots where each dot represents a gene in the (E) pink or (F) greenyellow module. Red and blue dots indicate up- or downregulation in PAH lungs, respectively. Genes are plotted by the Pearson correlation of their expression with PAH diagnosis (vs control) on the y-axis and the module eigengene on the x-axis. (G) Dot plot showing the normalized enrichment score (NES) of modules for the PAH lung differential transcriptome as determined by GSEA. Larger size dots indicate stronger FDR value. (H-I) Dot plots showing enrichment of modules for PAH GWAS SNPs using two distinct computational methods, (H) MAGMA, and (I) GSA-SNP2, across four independent PAH GWAS cohorts. Vertical red dotted lines indicate significance threshold. SNPs were mapped to genes by chromosomal proximity (within 20 kilobases from the 5’ or 3’ ends of a gene) and genes were scored for association with PAH based on disease-SNP p-value associations from GWAS summary statistics. Gene scores were then used in competitive gene-set analyses to identify module enrichment for PAH common genetic variation. To aggregate genetic variants into a gene score, the mean χ^2^ statistic and the log-minimum GWAS p-value for all SNPs localizing to a gene were used in MAGMA and GSA-SNP2, respectively. To determine significance, MAGMA uses a linear mixed model whereas GSA-SNP2 uses a standard normal distribution. Both methods adjust for gene size and gene density (the number of SNPs assigned to a given gene). WGCNA = weighted gene co-expression network analysis; GWAS = genome-wide association study; scRNAseq = single-cell RNA sequencing; Dx = diagnosis; PFT = pulmonary function test; RHC = right heart catheterization; Histo = histology; DLCO = diffusing capacity for carbon monoxide; FVC/DLCO = ratio of forced vital capacity to DLCO; mPAP = mean pulmonary artery pressure; PVR = pulmonary vascular resistance; REVEAL = Registry to Evaluate Early and Long-Term PAH Disease Management; cor = correlation; PAHB = US PAH Biobank; PHAAR = French Pulmonary Hypertension Allele-Associated Risk; BHFPAH = British Heart Foundation Pulmonary Arterial Hypertension; UK = UK National Institute for Health Research BioResource (NIHRBR); MAGMA = Multi-marker Analysis of GenoMic Annotation; FDR = false discovery rate.

### The pink module is enriched for PAH genetic variations

Having identified PAH-relevant modules, we next asked whether these modules might be a cause or consequence of PAH pathogenesis. To infer causality, we integrated PAH genetic association studies with our lung-derived modules. Specifically, we employed two distinct computational approaches, MAGMA(17) and GSA-SNP2(18), to test whether modules were enriched for PAH-associated single nucleotide polymorphisms (SNPs) using the full summary statistics from four independent PAH genome-wide association studies (GWAS) totaling 11,744 individuals(19, 20): the US PAH Biobank (PAHB), French Pulmonary Hypertension Allele-Associated Risk (PHAAR), British Heart Foundation Pulmonary Arterial Hypertension (BHFPAH), and UK National Institute for Health Research BioResource (NIHRBR) cohorts. Despite different statistical methods, MAGMA and GSA-SNP2 captured similar relative associations of genes with PAH genetic variation and neither approach was biased towards gene size or number of SNPs localizing to a gene (Supplemental Figure 4). We found that only the pink module was significantly enriched for PAH-associated SNPs using both approaches and across multiple cohorts (Figures 2H-2I): PAHB and PHAAR cohorts by both methods, and BHFPAH using GSA-SNP2. This finding suggests that pink module genes are not only associated with PAH diagnosis and severity, but also enriched with genetic risk of developing PAH.

### The pink module is co-regulated with known PAH genes and is enriched for Wnt signaling and EMT pathways

To delineate the regulatory relationships among genes within co-expression modules, we employed a Bayesian network analysis to build a gene regulatory network of the human lung by incorporating 2,066 lung transcriptomes, lung-specific expression quantitative trait loci (eQTL), and known transcription factor-target gene relationships (Supplemental Figure 5). Projection of co-expression module genes onto this lung regulatory network confirmed that the genes within individual modules are in close neighborhoods in the gene regulatory network analyses (Figure 3A).

**Figure 3:**
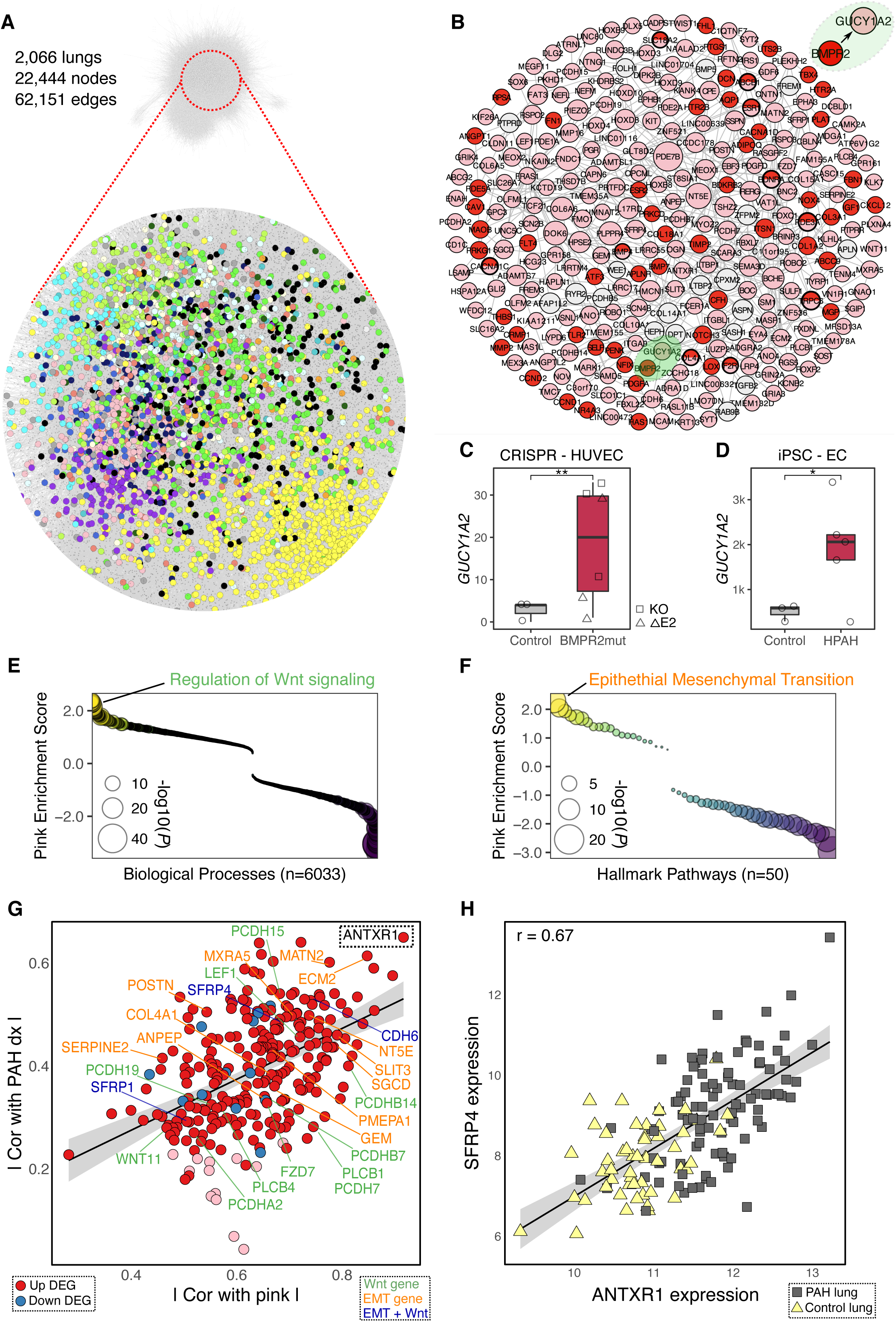
The pink module is co-regulated with known PAH genes and is enriched in Wnt signaling and EMT pathways. (A) Bayesian gene regulatory network constructed from 2,066 lung transcriptomes with incorporation of lung-specific expression quantitative trait loci (eQTL), and known transcription factor-target gene relationships. Nodes represent genes. Edges represent inferred gene-gene regulation. Node positions were determined by a prefuse force-directed algorithm. Genes from co-expression modules are shown projected onto this regulatory network with the different color nodes representing module membership. The largest modules with >3000 genes (turquoise, blue, and brown) are not shown to allow better visualization of other modules. (B) Pink subnetwork where pink genes and known PAH genes (red nodes) from disease-gene databases (Comparative Toxicogenomics Database(11) and DisGeNET(10)) were projected onto the lung Bayesian regulatory network in (A). BMPR2-GUCY1A2 pair is highlighted in the upper right where the arrow represents the predicted directional regulatory relationship. Larger size nodes represent hub genes where node size is proportional to -log10(FDR) as determined by Key Driver Analysis (4, 28, 29). Light grey nodes represent hub genes of the pink subnetwork that are neither pink nor red genes. (C-D) Box plots showing *GUCY1A2* expression two independent RNA-seq datasets: (C) CRISPR/Cas9-induced monoallelic mutations in BMPR2 (n = 6) vs wild-type control (n = 3) human umbilical vein endothelial cells (HUVEC)(30) and (D) endothelial cells (ECs) derived from induced pluripotent stem cells (iPSCs) of patients with hereditary PAH (HPAH) due to BMPR2 mutations (n = 5) vs control (n = 3)(31). *P* values were determined by DESeq2 for (C) and (D). (E-F) Dots plots showing Gene Set Enrichment Analysis (GSEA) of the pink module signature using (E) Gene Ontology (GO) and (F) Hallmark(13) gene sets where y-axis represents normalized enrichment scores (NES) in which scores greater than or less than zero represent gene sets enriched in genes positively or negatively correlated with the pink eigengene, respectively. The x-axis represents gene sets ordered by their enrichment score. Select top gene sets are labeled: Regulation of Wnt signaling (NES score 2.29, NES rank 4 of 6,033) in (E) and Epithelial Mesenchymal Transition (NES score 2.30, NES rank 1 of 50) in (F). Dots larger in size represent higher -log10(*P*) values. (G) Scatter plot showing pink genes where the x- and y-axes represent the absolute correlation of the pink gene with the pink eigengene and PAH diagnosis, respectively. Red and blue dots denote up- and downregulated genes in PAH lungs, respectively. EMT genes from Hallmark(13) and Wnt genes from PANTHER(69) are indicated by green and orange text, respectively, and blue text indicates genes in both gene sets. (H) Scatter plot showing expression of ANTXR1 (x-axis) vs. SFRP4 (y-axis) in PHBI lungs where yellow triangles and grey squares represent control and PAH lungs, respectively. * *P* < 0.05, ** *P* < 0.01. BMPR2mut = BMPR2 mutation; KO = knockout; rE2 = deletion in exon 2; CRISPR = clustered regularly interspaced short palindromic repeats.

We found that a number of established PAH genes co-localize with pink module genes in the Bayesian gene regulatory network, suggesting a regulatory relationship between PAH genes and pink module genes (Figure 3B). For example, BMPR2, the most well-established causal PAH gene, was predicted to regulate GUCY1A2, a pink module gene that is also upregulated in PAH lungs (Supplemental Figure 6). To validate this prediction, we queried public RNA-seq datasets (30, 31) and found that *GUCY1A2* was upregulated in CRISPR-induced BMPR2 mutant endothelial cells (ECs), and in ECs derived from induced pluripotent stem cells (iPSCs) from hereditary PAH patients with BMPR2 mutations (Figures 3C-3D).

We then used gene set enrichment analysis (GSEA) to functionally characterize the pink signature which we defined as the correlation of the pink eigengene with the expression of genes across the transcriptome. We found that regulation of Wnt signaling and epithelial mesenchymal transition (EMT), both important pathways in PAH(48–50), were strongly enriched in the pink module (Figures 3E-G). *ANTXR1* was the top pink hub gene most connected to pink module genes and most correlated with PAH (Figure 3G). While not traditionally associated with Wnt signaling or EMT, *ANTXR1* has been recently implicated in various cancers through such pathways(51, 52). Furthermore, the strongest pairwise connection among pink genes as determined by WGCNA (out of >30k pairs) was between *ANTXR1* and *SFRP4*, a secreted frizzled-related protein and one of 3 pink genes known to be involved in both Wnt signaling and EMT (Figures 3G-3H). Therefore, the pink module and its top PAH-associated gene *ANTXR1* may play a role in PAH through modulation of Wnt signaling and EMT.

### Cell type deconvolution reveals cell-type specificity in PAH lung modules

Having identified PAH-specific transcriptional changes at the whole lung level, we next asked whether cell-type fractional changes could be inferred from the transcriptomes of PAH and control lungs by deconvolution analysis based on transcriptomic references of 37 lung cell type clusters from seven publicly available human lung single-cell RNAseq datasets (34–40) (Figures 4A-4B; Supplemental Figure 7). Using the cell-type references from this integrated reference atlas, we deconvoluted PHBI bulk transcriptomes using CIBERSORTx(41) and found that PAH samples clustered together based on estimated cell fractions (Supplemental Figure 8) and that specific cell-type fractions correlated positively or negatively with PAH, such as endothelial (i.e. lymphatic and arterial) and myeloid (i.e. interstitial macrophage and classical monocyte) subpopulations, respectively (Figure 4C). In addition to lymphatic and arterial endothelial cells, myofibroblast fractions were particularly abundant in PAH samples relative to control, whereas cell fractions of other vascular mesenchymal subpopulations (i.e. fibroblast, smooth muscle cell and pericyte) were unchanged. Endothelial capillary 1 (EndoCap1) fractions were decreased in PAH lungs consistent with microvascular rarefaction, a known feature of PAH(53).

**Figure 4:**
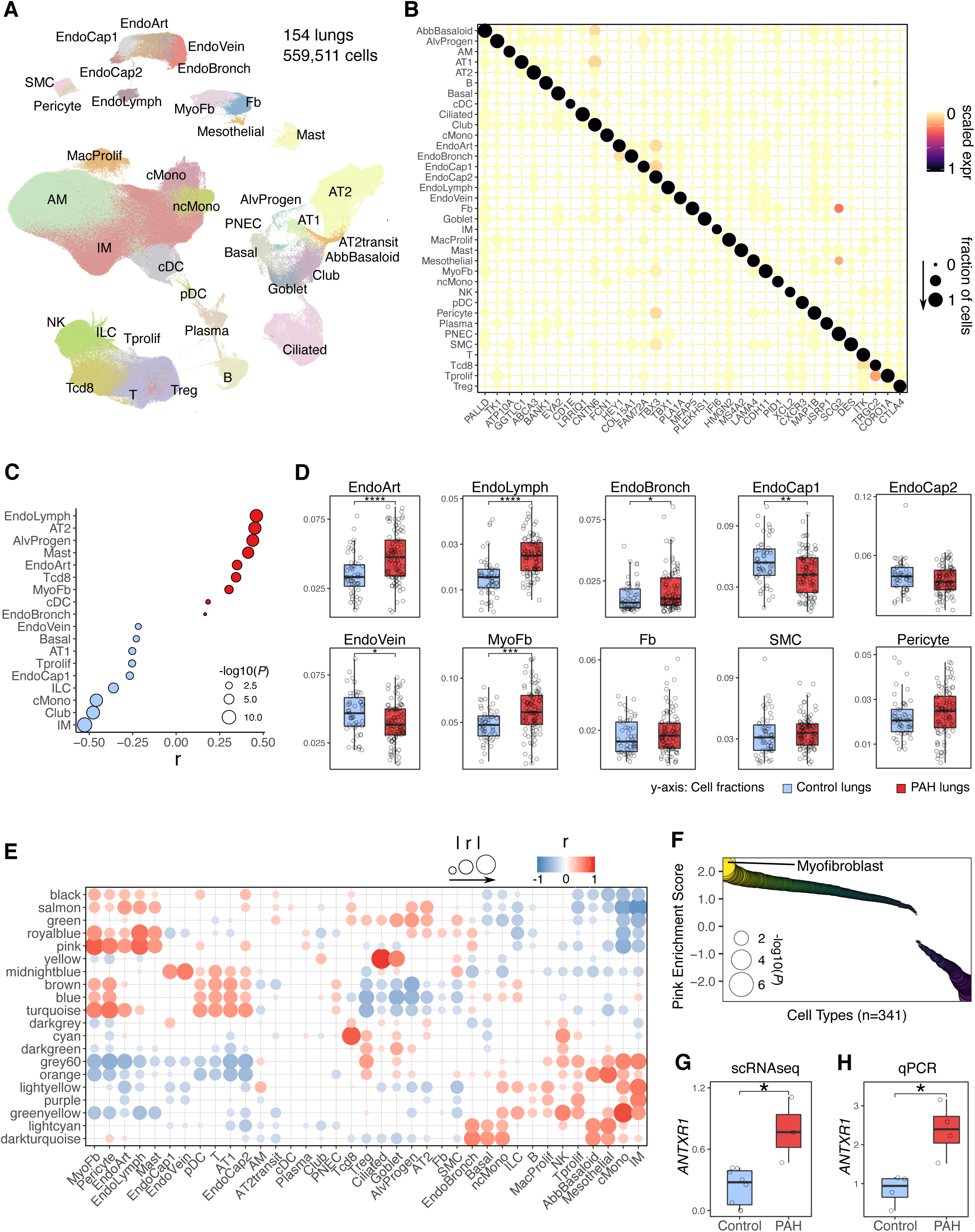
Deconvolution reveals cell-type specificity in PAH lungs and modules. (A) Uniform Manifold Approximation and Projection plot showing seven publicly available human lung single-cell RNAseq datasets(34–40) integrated and reclustered in Seurat(70), totaling 559,511 cells from 154 lungs. (B) Heatmap showing scaled expression of cell-type specific marker genes on the x-axis across all cell types in (A) on the y-axis. Larger size dots indicate higher fraction of cells expressing a given gene. (C) Dot plot showing Pearson correlation of deconvoluted cell type fractions with PAH vs control status across PHBI lung samples. Only correlations with *P* < 0.05 are shown. (D) Box plots of deconvoluted vascular cell fractions in PAH vs control lungs. (E) Heatmap showing Pearson correlation of deconvoluted cell fractions (columns) and module eigengenes (rows). Only correlations with *P* < 0.05 are shown. Larger size dots indicate higher absolute correlation. (F) Dot plot showing the normalized enrichment score of the pink module signature (defined as the correlation of the pink eigengene with the expression of genes across the transcriptome) for known cell-type signatures from Azimuth(15) as determined by GSEA. Larger size dots indicate stronger *P* value. (G) Box plot showing averaged *ANTXR1* expression in fibroblasts profiled by scRNAseq from 3 PAH vs 6 control lungs (32). Note, myofibroblasts were not subclustered in this dataset. (H) Box plot showing *ANTXR1* expression by qPCR of pulmonary arterial adventitial cells isolated from PAH vs. control lungs (n = 4 biological replicates/group). Wilcoxon rank-sum test: * *P* < 0.05, ** *P* < 0.01, *** *P* < 0.001, **** *P* < 0.0001. AbbBasaloid = aberrant basaloid; AlvProgen = alveolar progenitor; AM = alveolar macrophage; AT1 = alveolar type 1; AT2 = alveolar type 2, cDC = conventional dendritic cell; cMono = classical monocyte; EndoArt = endothelial arterial; EndoBronch = endothelial bronchial; EndoCap1 = endothelial capillary 1; EndoCap2 = endothelial capillary 2; EndoLymph = endothelial lymphatic; EndoVein = endothelial vein; Fb = fibroblast; IM = interstitial macrophage; MacProlif = macrophage proliferating; MyoFb = myofibroblast; ncMono = nonclassical monocyte; NK = natural killer; pDC = plasmacytoid dendritic cell; PNEC = pulmonary neuroendocrine cell; SMC = smooth muscle cell; Tcd8 = CD8^+^ T cell; Tprolif = proliferating T cell; Treg = regulatory T cell; scRNAseq = single-cell RNA sequencing; qPCR = quantitative polymerase chain reaction.

We then integrated the deconvoluted cell fractions with our bulk lung-derived co-expression modules to decipher cell-type specificity of individual modules. We identified distinct patterns of correlation between cell fractions and module eigengenes where, for example, some modules were highly specific to particular cell types such as yellow to ciliated cells and cyan to CD8^+^ T cells (Figure 4E). Among the top correlations was the PAH-associated pink module to myofibroblast fractions (r = 0.78). In a complementary approach to infer cell specificity, we then performed GSEA using known cell-type signatures of 341 cell types across >9 tissues and found the pink module to be most enriched in the myofibroblast signature (Figure 4F). Given this finding, we then asked whether *ANTXR1*, the top pink gene whose expression is upregulated in PAH lungs, might be upregulated in PAH lung fibroblasts specifically. Supporting our cell-type deconvolution analyses, we found that *ANTXR1* is upregulated in PAH fibroblasts by lung single-cell RNA-sequencing in a published dataset (in which myofibroblasts were not subclustered) (32) and in pulmonary arterial adventitial cells isolated from PAH lungs by qPCR (Figures 4G-4H).

### Pharmacotranscriptomics identifies novel therapeutic targets

Having prioritized the pink module by association with PAH diagnosis, clinicopathologic severity, and genetic risk, we next investigated whether pattern matching the pink module signature with known pharmacologic and genetic perturbation signatures could reveal novel therapeutic targets. We screened the pink signature against 8,559 perturbation signatures from Connectivity Map (CMap) (42). These CMap signatures were grouped into 171 classes that share similar mechanisms of action or biological functions. We found ubiquitin specific peptidase (USP) loss-of-function by short hairpin RNAs (shRNAs) to have the highest connectivity score of all CMap classes, which suggests that knockdown of USPs induces a transcriptional response highly similar to the pink signature (Figure 5A). Indeed, six different USPs (USP7, USP22, USP12, USP20, USP1, and USP15) were among the top scoring perturbations (Figure 5B). We next queried an independent genetic perturbation screen of 7,502 genes by CRISPR knockout (KO) from the LINCS L1000 database(44) and found that USP28 KO was a top mimicker of the pink signature (Figures 5C-5D). Therefore, targeting members of the USP family by either complete knockout via CRISPR or partial knockdown via shRNA induced transcriptional changes similar to that of pink module genes in PAH lungs.

**Figure 5:**
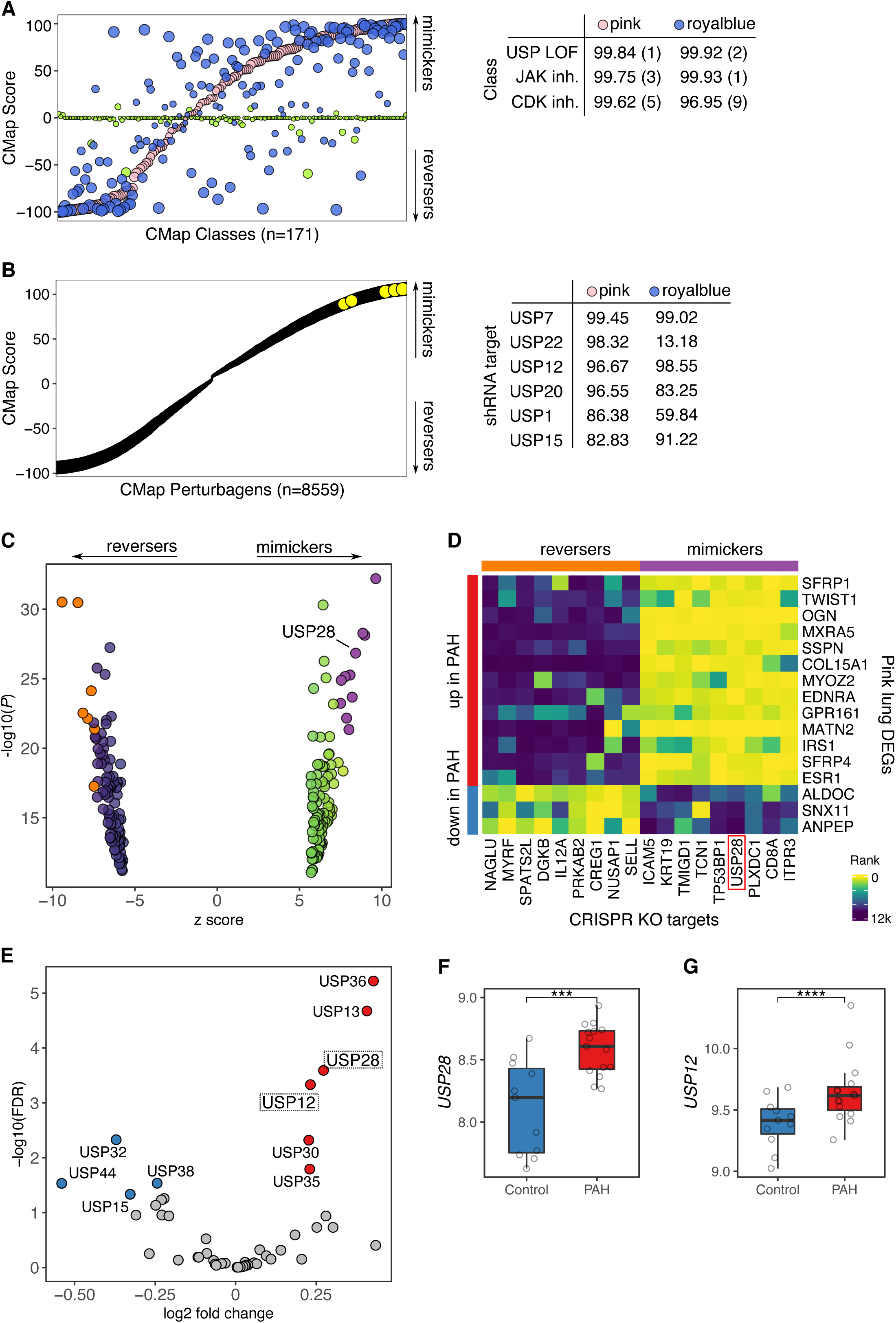
Pharmacotranscriptomics identifies novel therapeutic targets. (A) Dot plot showing CMap score of 171 CMap classes which are groups of pharmacologic or genetic perturbagens that share the same mechanism of action or biological function. Scores approaching 100 or -100 indicate perturbagens predicted to mimic or reverse the query signature, respectively. Larger size dots indicate higher absolute score. Color indicates module membership. Table on the right shows the score of select CMap classes with the rank out of 171 classes shown in parenthesis. (B) Dot plot showing CMap score of 8,559 CMap perturbagens. Table on the right shows CMap scores of USP shRNA targets. (C) Volcano plot of top 100 mimickers and top 100 reversers of the pink PAH DEG signature out of a total of 7,502 genetic targets screened from the CRISPR KO consensus signature database of LINCS L1000 using SigCom(44). Z score on the x-axis indicates the degree to which the target signature mimics or reverses the pink signature. Select top mimickers and reversers that are shown in a heatmap in (D) are labeled by purple and orange, respectively. (D) Heatmap showing the relative expression of select pink lung DEGs (rows) in the signatures of select top CRISPR KO targets (columns) that are mimickers or reversers of the pink signature. Lowly ranked genes are downregulated while highly ranked genes are upregulated in the CRISPR KO vs. control. Rows are annotated on the left by red or blue to indicate upregulated or downregulated in PAH vs. control lungs, respectively. (E) Volcano plot showing upregulated and downregulated USPs in PAH lungs in red and blue, respectively. Grey dots indicate genes with FDR ≥ 0.05. USPs that were both upregulated and top hits in our CMap or CRISPR analyses are boxed. (F-G) Box plots showing mRNA expression of (F) USP28 and (G) USP12 in a human whole lung microarray of 15 PAH patients vs 11 controls(33). *P* values were obtained from NCBI’s GEO2R: \*\*\**P* < 0.001, **** *P* < 0.0001. CMap = Connectivity Map; USP = ubiquitin specific peptidase; JAK inh. = Janus kinase inhibitor; CDK inh. = cyclin-dependent kinase inhibitor; shRNA = short hairpin RNA.

Interestingly, Janus kinase (JAK) and cyclin-dependent kinase (CDK) inhibitors, both recently studied as potential therapies in PAH(54–56), were also among the top CMap classes whose transcriptional signature matched that of the pink module (Figure 5A). We found a similar connectivity profile for another co-expression module, royalblue, containing a distinct set of 98 genes which also shared with pink a similar pattern of clinicopathologic correlations (Figures 5A and 2D) and was second only to pink as a top enriched module for upregulated PAH DEGs (Figure 2G). This suggests converging and targetable pathways between pink and royalblue genes. In contrast, greenyellow, the most enriched module for downregulated DEGs, did not show a strong CMap matching profile (Figure 5A). Indeed, royalblue was also enriched for Wnt signaling and EMT and a number of its module member genes co-localized to the pink sub-regulatory network (Supplemental Figure 9) such as LTBP2, a recently identified biomarker for PAH(57).

Given that JAK and CDK inhibitors, both of which mimicked the pink signature (Figure 5A), may have a therapeutic role in PAH based on recent preclinical studies(54–56) and given that the pink eigengene correlated with clinical markers of favorable disease prognosis (Figure 2D), we reasoned that the pink signature could be beneficial in PAH. As downregulation of USPs induced transcriptomic signatures that mimicked that of the pink module in our CMap and L1000 analyses, we postulate that targeting USPs might also have a therapeutic role. We found that USP28 and USP12, whose downregulation led to transcriptomic responses mimicking that of the pink module, were upregulated in PAH lungs in our PHBI dataset (Figure 5E) and in an independent microarray (Figures 5F-5G). These USPs may serve as top candidates for PAH therapeutic development.

## Discussion

Leveraging the largest PAH lung biobank to date and the first to use RNA-sequencing combined with state-of-the-art multiomic integration and systems biology approaches, we dissected the transcriptional landscape of PAH lungs to uncover a novel gene module enriched in upregulated genes and associated with clinicopathologic severity, genetic risk, specific vascular cell types, and new therapeutic targets in PAH.

We identified by network analysis a set of 266 co-expressed genes called the pink module that was not only associated with objective measures of underlying disease severity such as increased PVR, increased intimal thickness, and reduced DLCO, but also associated with lower risk of mortality by REVEAL lite as well as indicators of clinically compensated PAH such as lower number of PAH hospitalizations, signs of right heart failure, WHO functional class and NT-proBNP (Figure 2D). We hypothesize that the pink module is active in response to the underlying disease process to counteract disease progression in PAH. Supporting this possibility, JAK and CDK inhibitors, both of which counteract preclinical PAH (54–56), were top perturbagens predicted to mimic the pink signature in our CMap analysis (Figure 5A). Moreover, our regulatory network analysis uncovered a novel connection where deficiency of BMPR2, the most well-established causal PAH gene, leads to an upregulation of pink gene GUCY1A2 (Figures 3B-3D). GUCY1A2 encodes the alpha subunit of soluble guanylate cyclase 1 (GC-1), the primary receptor of nitric oxide and the stimulation of which is the primary mechanism of action of riociguat, an FDA-approved therapy in clinical use to treat PAH patients. Therefore, supporting our hypothesis that the pink module might be a response to PAH to counteract the disease, deficiency of BMPR2, which is causative and harmful in PAH pathogenesis, leads to upregulation of the pink module gene GUCY1A2 which is beneficial in PAH.

EMT and Wnt signaling were top pathways enriched in the pink module, both of which are known to play a critical role in PAH pathobiology and are interrelated (Wnt signaling induces EMT) (Figures 3E-3G)(48, 49, 58, 59). In terms of specific pink genes and their potential role in these pathways, ANTXR1, a transmembrane protein that interacts with extracellular matrix proteins, was the top hub gene most connected to other pink module genes and it was upregulated in not only PAH lungs (Figure 1C) but specifically in PAH lung fibroblasts (Figures 4G-4H). While its role in PAH has not been investigated, one study found that ANTXR1-deficient fibroblasts showed increased expression of EMT markers *Col1a1* and *Fn* raising the possibility that ANXTXR1 might modulate EMT in a beneficial manner in PAH(60). Furthermore, SFRP4, which showed the strongest pairwise correlation with ANTXR1 among all pink gene pairs (Figure 3H), is a secreted Wnt antagonist that has been shown to inhibit EMT in cancer cells(61, 62). Therefore, the pink module may counteract the PAH disease process by modulation of EMT and Wnt signaling.

The following question then arises-if the pink module and its member genes (i.e. ANTXR1, SFRP4, and GUCY1A2) are beneficial in PAH and upregulated in explant lungs, why did the disease in these patients still progress to the point of needing a lung transplantation? One possibility is that the pink module is activated too little and/or too late in these patients or the activation is not sufficient to counteract the effects of other deleterious pathways. Despite its relatively large size of 266 genes, the pink signature can be leveraged using pharmacotranscriptomic pattern matching algorithms to identify novel therapeutic targets for testing in future investigations with preclinical models. Using such an approach, we identified JAK and CDK inhibitors as well as ubiquitin specific peptidase (USP) loss-of-function as CMap perturbagen classes which induce signatures that match that of pink, but also ubiquitin specific peptidase (USP) loss-of-function as the top CMap class mimicker of the pink signature. As main members of the deubiquitinase family, USPs are involved in diverse processes such as cell cycle progression, apoptosis, EMT, and DNA damage repair and have been strongly implicated in cancer progression(63). Furthermore, USPs regulate PAH-relevant pathways such as NFkB, TGFbeta, and Wnt signaling and have been investigated as therapeutic targets in cancer and other fields(64, 65), yet their role in PAH has not yet been described. We demonstrated that targeting members of the USP family by either complete knockout via CRISPR or partial knockdown via shRNA induced transcriptional changes similar to that of pink DEGs in PAH lungs (Figure 5). Specifically, USP28 and USP12 may serve as particularly attractive targets for downstream investigation as they were also upregulated in PAH lungs (Figures 5E-5G). While studies on USP12 are limited, USP28 has been shown to activate Wnt signaling(66, 67) and its inhibition blocks EMT progression in cancer cells(68). Thus, inhibition of Wnt signaling and EMT may be the common pathways shared between USP loss-of-function and the pink module.

Given that the PHBI cohort consisted of patients with advanced stage PAH at the time of lung transplantation, our results are likely not representative of the full range of disease and our analysis was limited in discerning cause versus consequence of PAH. However, the deep clinical phenotyping allowed us to make correlations with disease severity, and GWAS integration enabled us to infer causality in PAH pathogenesis. The majority of patients had idiopathic PAH and thus our findings may not be generalizable to other WHO Group 1 PAH subtypes or other WHO groups, and our sample size of other PAH subtypes was insufficiently powered to detect subtype-specific differences. The majority of our patients were also female, reflective of the strong female predominance of PAH. However, our sensitivity analyses did not reveal significant sex-specific differences in the top pathway and module enrichments (Supplemental Figure 10). Finally, while heterogeneity of PAH-targeted therapy in these patients could affect the transcriptome profiles, the majority of patients were on triple therapy including prostacyclin infusion. Thus, we did not explore treatment-specific differences.

In conclusion, our study leverages the largest PAH lung biobank to date to provide an in-depth analysis of the lung transcriptional landscape of PAH using multiomic integration and systems biology approaches. Through this analysis, we uncovered a novel gene network module that is associated with PAH risk and severity, may counteract disease progression through modulation of EMT and Wnt signaling, and may be regulated by USPs. Future experimental studies such as knockdown of USPs in PAH vascular cells are warranted to further investigate the role and therapeutic potential of the pink module and targeting USPs in PAH.

## Acknowledgements

The authors thank the investigators, personnel, and participants of the PHBI, particularly those involved in the transplant and preparation centers: Allegheny University of the Health Sciences (PI: Raymond L. Benza, M.D.); Baylor College of Medicine (George Noon, M.D.); Cleveland Clinic (PI: Serpil Erzurum, M.D.); Duke University (PI: Pang-Chieh Jerry Eu, M.D.); Stanford University-UCSF (PI: Marlene Rabinovitch, M.D.); University of Alabama (PI: Keith Wille, M.D.; prior PI: Raymond L. Benza, M.D.); University of California, San Diego (PI: Patricia Thistlethwaite, M.D., Ph.D); Vanderbilt University (Barbara Meyrick, Ph.D.). Supported by ALA CA-675591 (J.H.) and NHLBI R01HL162124 (M.E.).

## Supplemental Figure

**Supplemental Figure 1:**
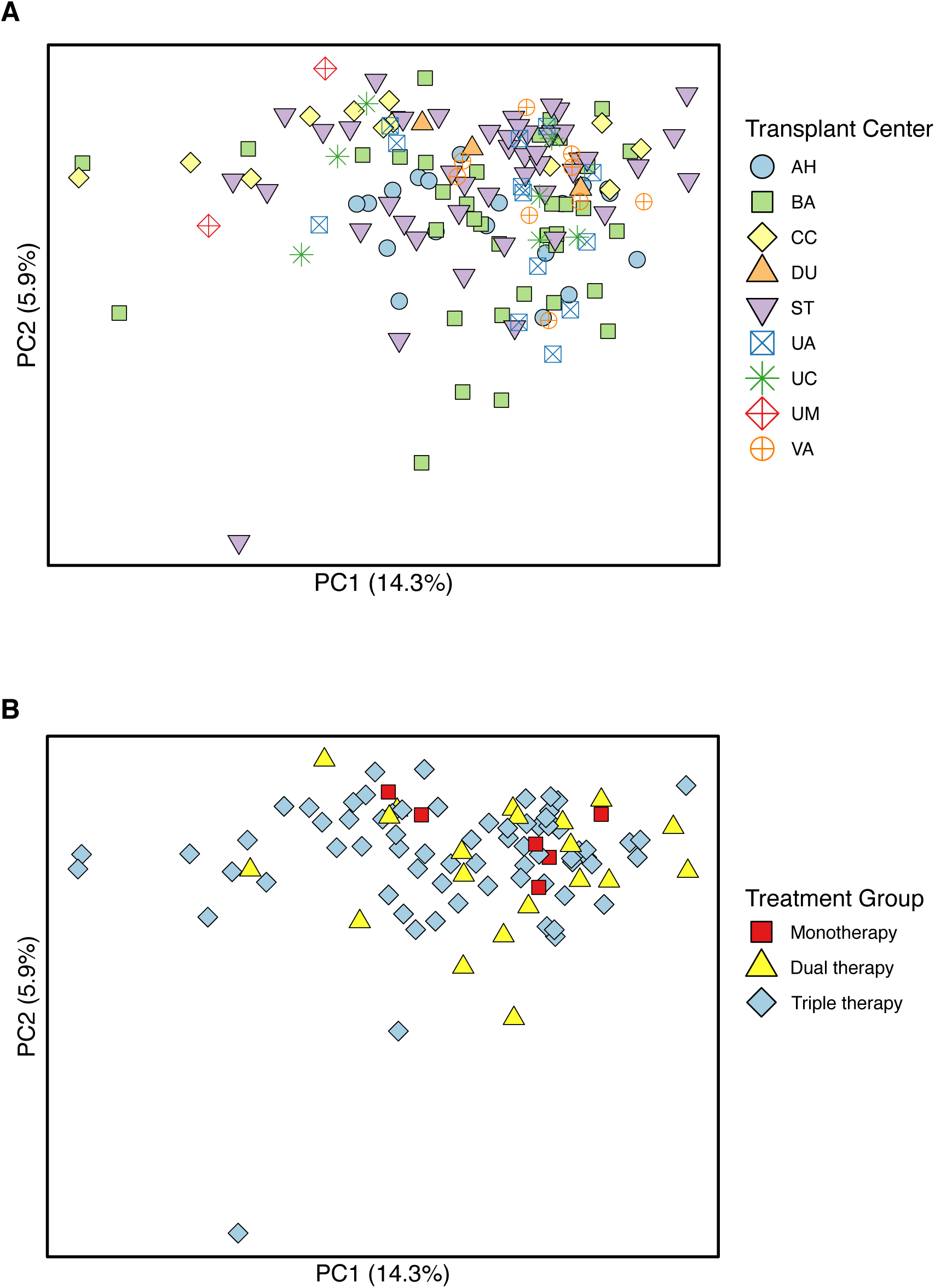
Lung transcriptomes do not cluster by transplant center nor treatment group. (A-B) PCA plots showing PCA of all 23,355 detected genes where samples are colored by (A) transplant center of tissue origin and (B) treatment group. Control samples are not shown in (B). Treatment groups refer to one or a combination of the three major classes of PAH-targeted drugs: phosphodiesterase type 5 inhibitors, endothelin receptor antagonists, and prostacyclin analogues. AH = Allegheny Hospital; BA = Baylor; CC= Cleveland Clinic, DU=Duke University; ST=Stanford; UA=University of Alabama; UC=University of California, San Diego; UM=University of Michigan; VA= Vanderbilt.

**Supplemental Figure 2:**
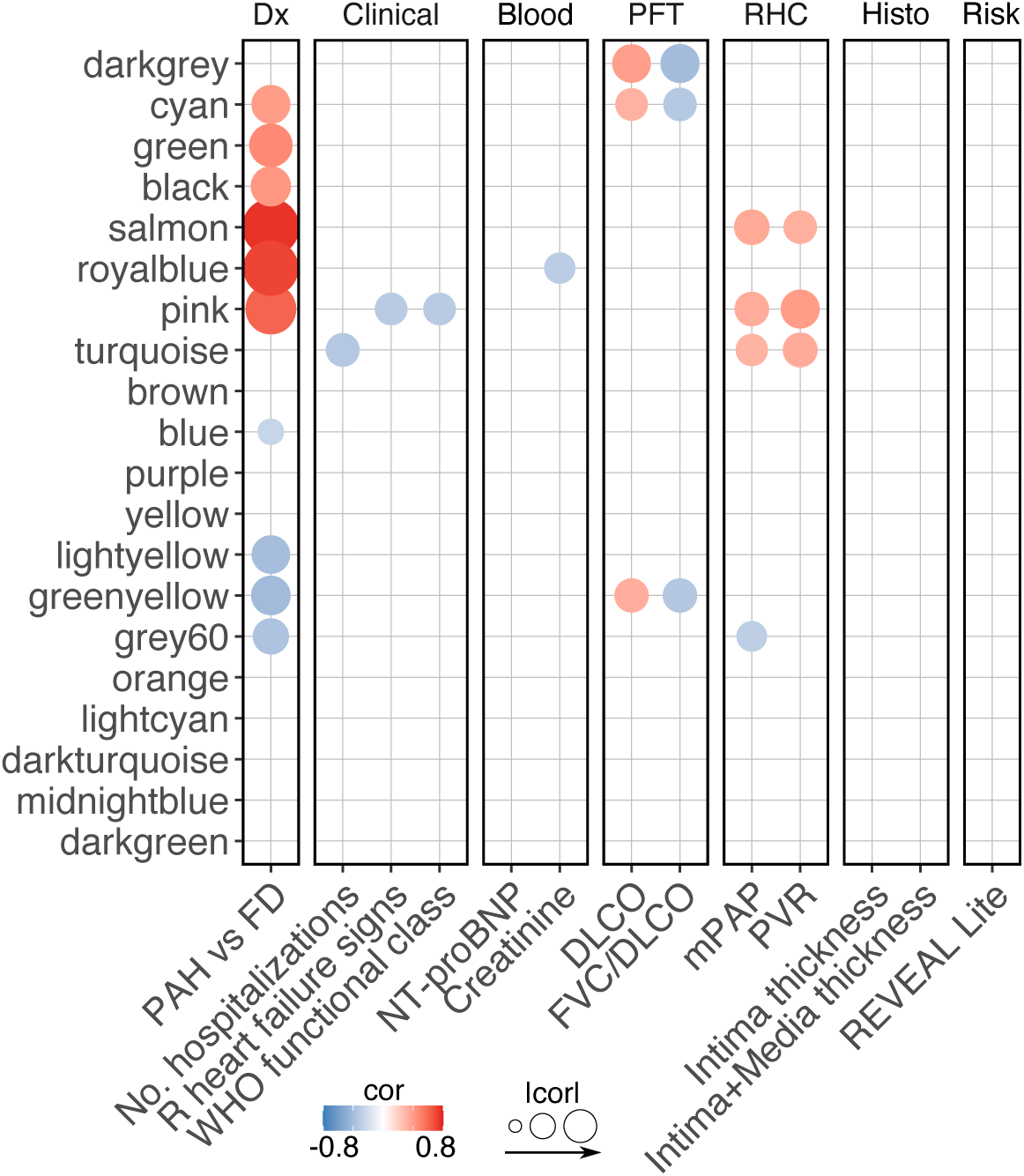
Modules correlated with clinical traits after adjustment for multiple testing. Heatmap showing Pearson correlations of module eigengenes with clinical and pathologic characteristics that met FDR threshold of < 0.05 after adjustment for multiple comparisons (n=260 comparisons from 13 traits and 20 modules). Red and blue dots indicate positive and negative correlation, respectively. Larger size dots indicate stronger correlation. No. hospitalizations indicates number of hospitalizations due to PAH between the time of diagnostic RHC and lung transplantation. R heart failure signs indicate signs of right heart failure such as ascites or leg swelling. Intima and intima plus media thickness were determined by morphometric analysis of volume density of pulmonary arteries in histological lung sections. Dx = diagnosis; PFT = pulmonary function test; RHC = right heart catheterization; Histo = histology; DLCO = diffusing capacity for carbon monoxide; FVC/DLCO = ratio of forced vital capacity to DLCO; mPAP = mean pulmonary artery pressure; PVR = pulmonary vascular resistance; REVEAL = Registry to Evaluate Early and Long-Term PAH Disease Management; cor = correlation.

**Supplemental Figure 3:**
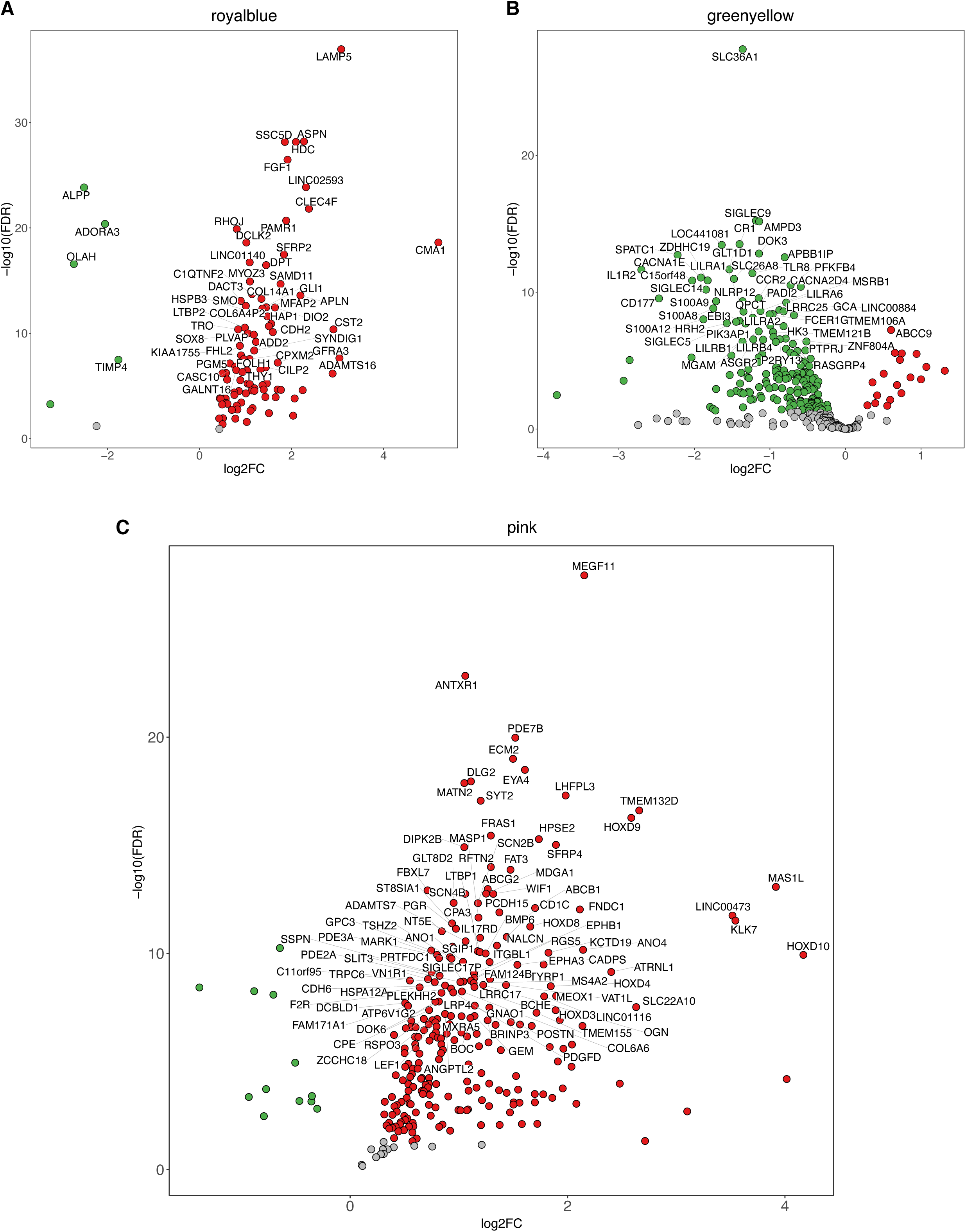
Dysregulated genes of royalblue, greenyellow and pink modules. (A-C) Volcano plots showing dysregulated genes of (A) royalblue, (B) greenyellow, and (C) pink where red, green, and grey indicate upregulation, downregulation, or no change in PAH, respectively. The top 50 (A-B) or 100 genes (C) by the absolute Wald statistic as determined by DESeq2 are labeled. FDR = false discovery rate; log2FC = log2 fold change.

**Supplemental Figure 4:**
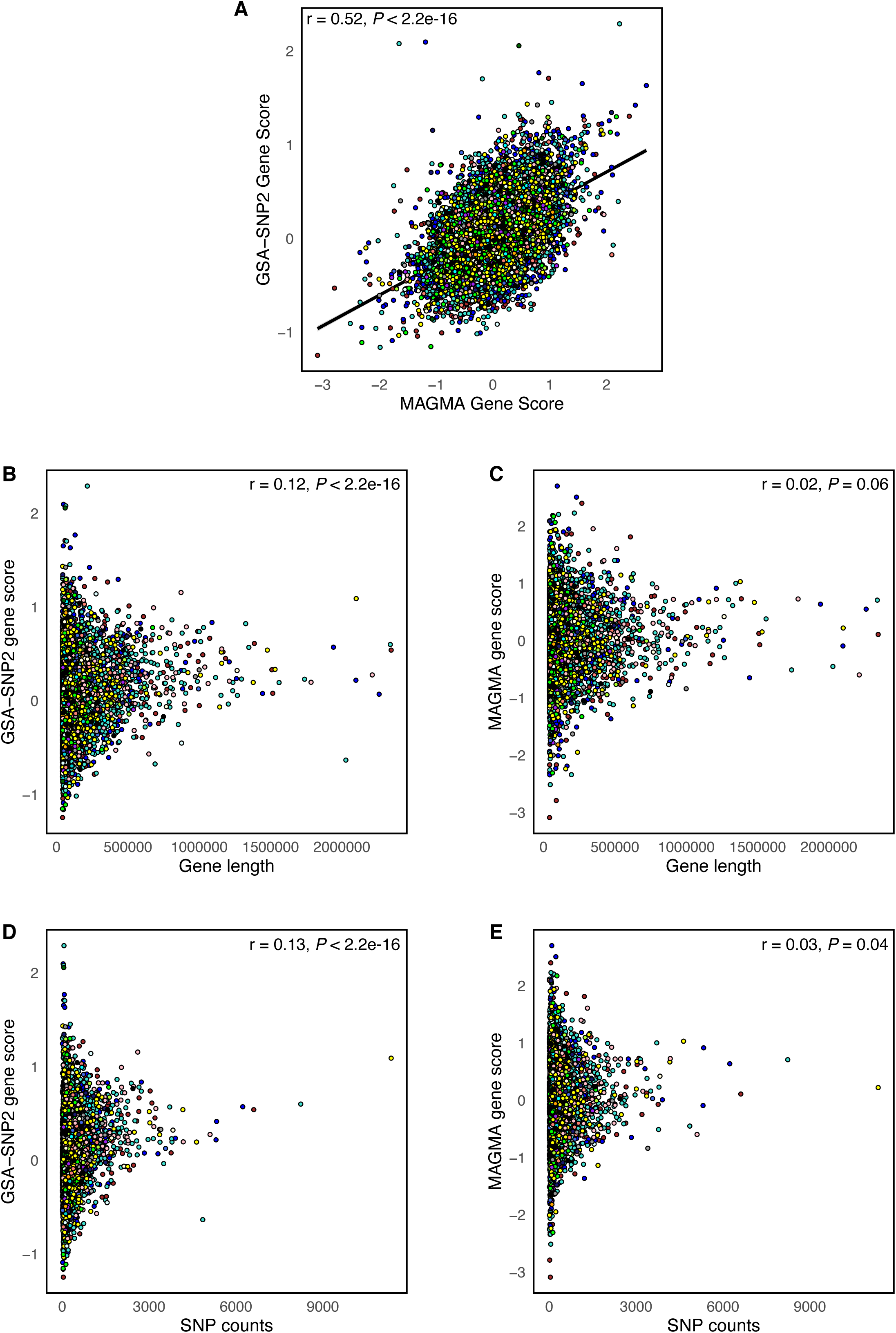
Analysis of MAGMA and GSA-SNP2 gene scores. (A) Scatter plot showing gene scores averaged across four PAH GWAS cohorts as determined by MAGMA on the x-axis and GSA-SNP2 on the y-axis. To aggregate genetic variants into a gene score, the mean χ^2^ statistic and the log-minimum GWAS p-value for all SNPs localizing to a gene were used in MAGMA and GSA-SNP2, respectively. (B-C) Scatter plots showing gene scores from (B) GSA-SNP2 and (C) MAGMA plotted against gene length in base pairs. (D-E) Scatter plots showing gene scores from (D) GSA-SNP2 and (E) MAGMA plotted against the average SNP counts across four PAH GWAS cohorts. SNP counts represent the number of SNPs localizing to a given gene within 20 kilobases from the 5’ or 3’ ends of the gene. (A-E) Colors represent module membership.

**Supplemental Figure 5:**
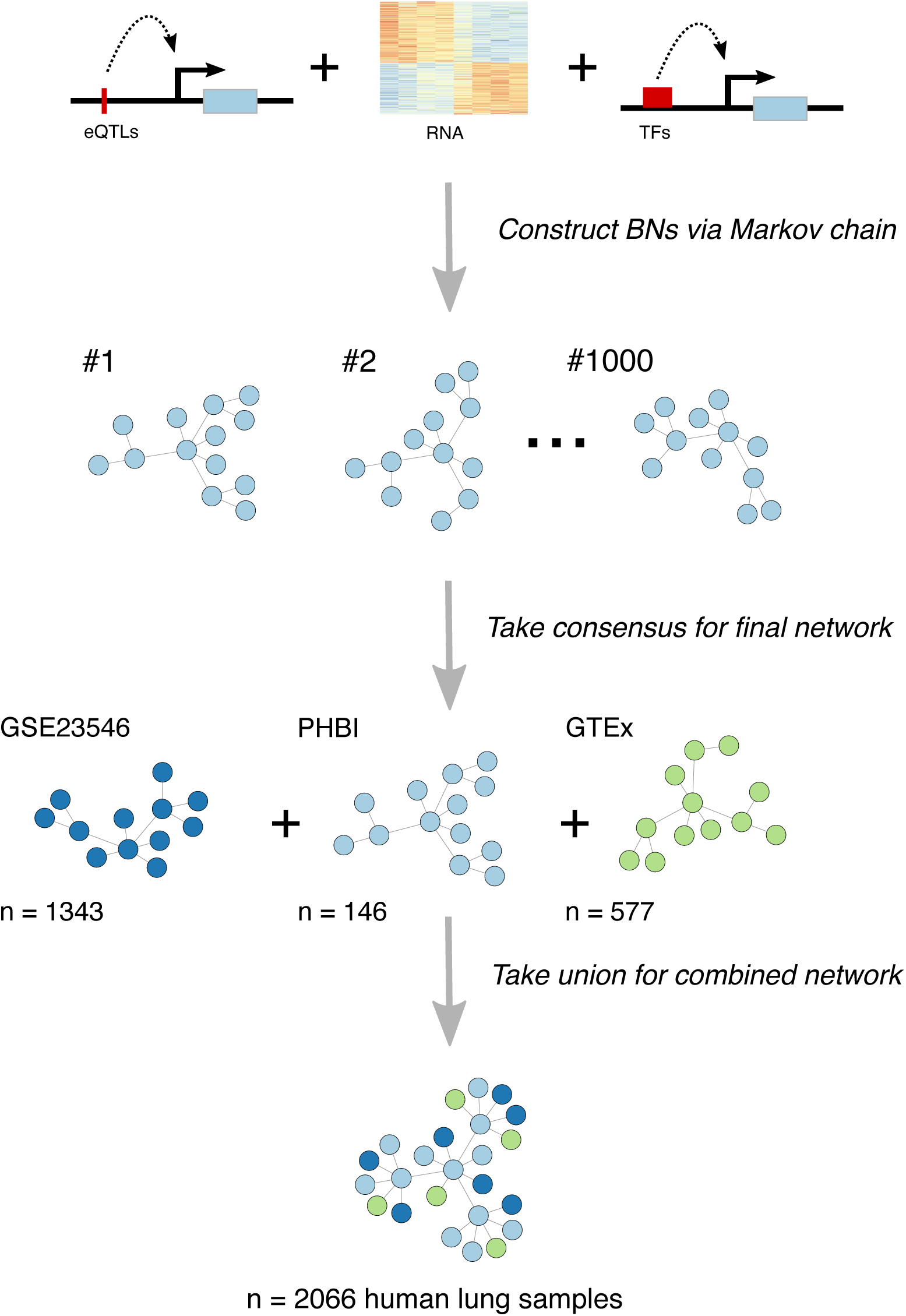
Bayesian network analysis workflow to construct a gene regulatory network of the human lung. Bayesian networks (BNs) were constructed using Reconstructing Integrative Molecular Bayesian Network (RIMBANet)^19^. For this method, 1000 networks were generated from different random seed genes using continuous and discrete expression data derived from transcriptomes from either GSE23546 (n = 1343) (1), PHBI (n = 146), or GTEx v8 (n = 577) (1). Whole lung-specific cis eQTLs from GTEx v8 (1) and transcription factor-target gene data from HTRI (1), TRRUST (1), and PAZAR (1) databases were used as priors. Then, the final network for each of the 3 datasets was obtained by taking a consensus network from the 1000 randomly generated networks whereby only edges that passed a probability of >30% across the 1000 BNs were kept. Finally, the union of the 3 networks was taken to create a combined gene regulatory network derived from a total of 2,066 human lungs.

**Supplemental Figure 6:**
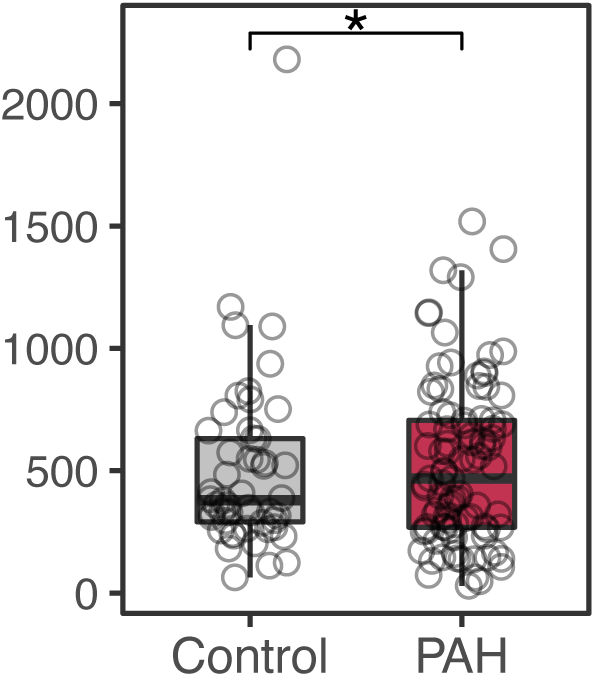
*GUCY1A2* is upregulated in PAH lungs. Box plot showing *GUCY1A2* expression in WHO Group 1 PAH (n = 93) vs control (n = 51) lungs from PHBI. * FDR < 0.05.

**Supplemental Figure 7:**
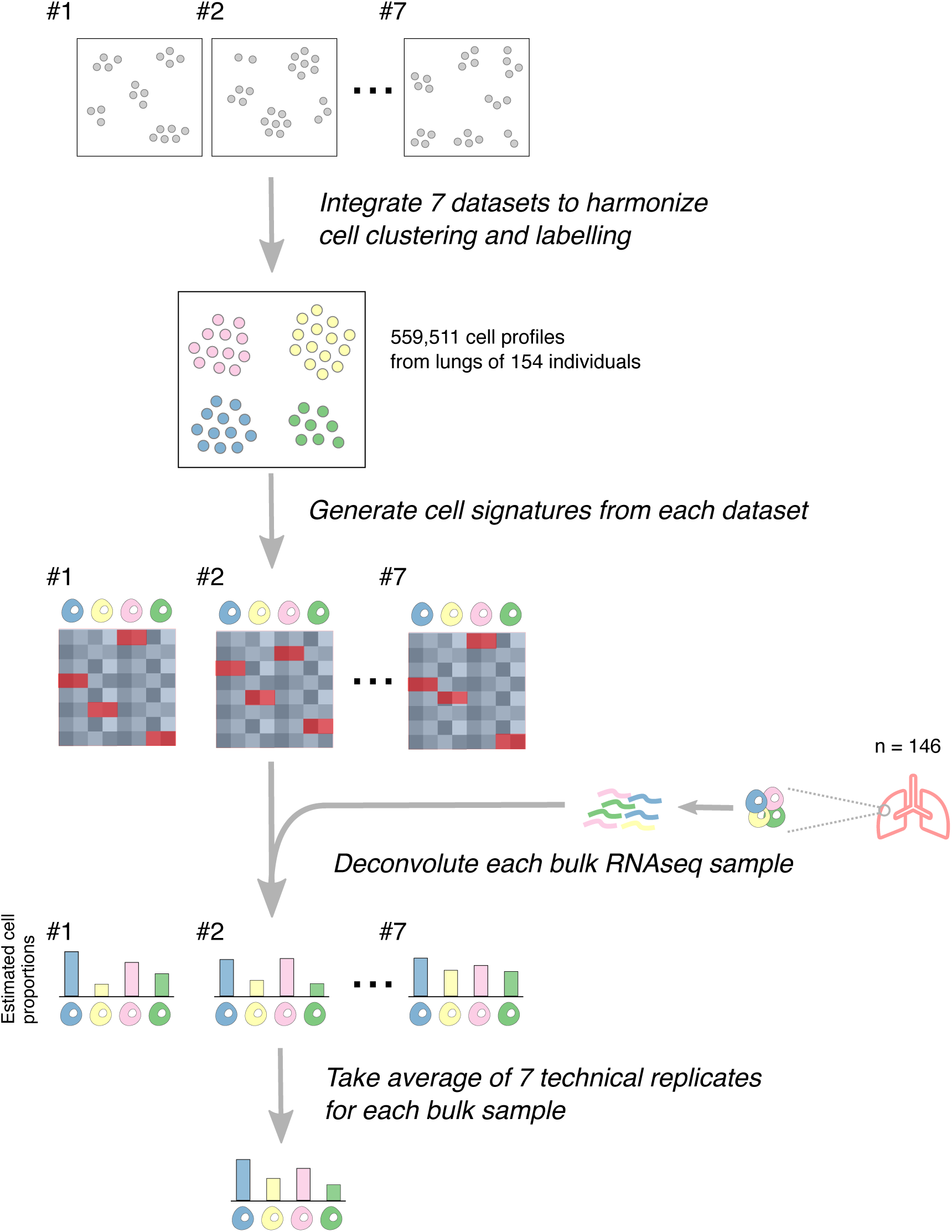
Single-cell RNA sequencing and deconvolution analysis scheme. To serve as a cell type reference for deconvolution, we integrated seven publicly available human lung single-cell RNAseq datasets(29–35) and identified 37 cell-type clusters using known marker genes from the literature. Within each cell-type cluster, the average expression of gene counts were calculated across cells for each individual sample to create a cell-type signature for each of the seven datasets. PHBI bulk transcriptomes were deconvoluted with CIBERSORTx(36) with cell-type signatures from each of the seven datasets as a reference. The resulting cell fractions using each of the seven dataset-specific reference signatures served as technical replicates. These technical replicates were then averaged to determine the final estimated cell fractions for each lung sample.

**Supplemental Figure 8:**
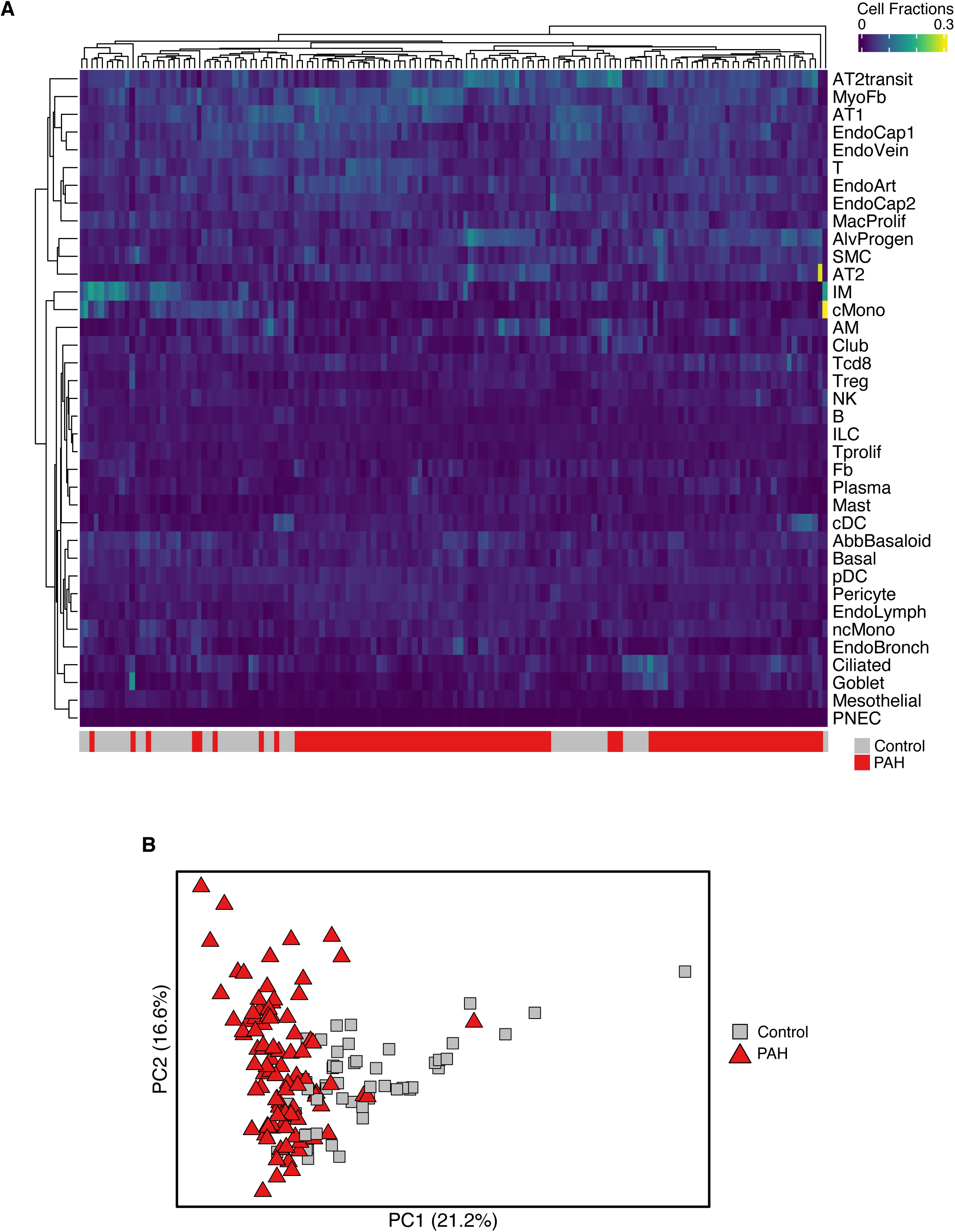
PAH lung samples cluster together based on estimated cell fractions. (A) Heatmap showing cell fractions estimated by deconvolution of PHBI lung transcriptomes by CIBERSORTx(1). Dendrograms are shown on the left and top representing hierarchical clustering of cell types (rows) and lung samples (columns), respectively. Lung samples are annotated at the bottom to indicate PAH in red or control in grey. (B) PCA plot showing PCA of estimated cell fractions with samples colored to indicate PAH in red or control in grey.

**Supplemental Figure 9:**
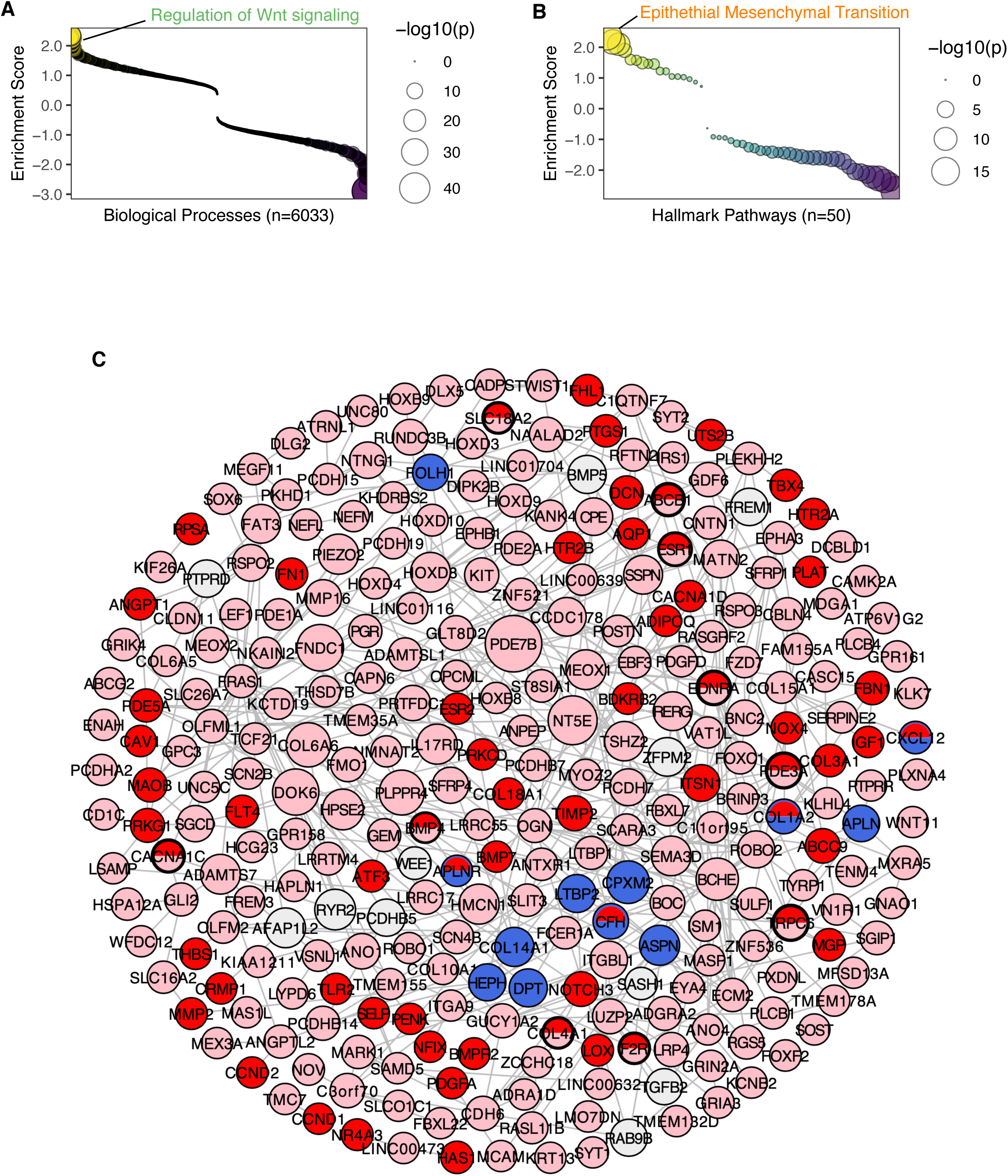
Royalblue genes share similar pathways with and are hub nodes of the pink module. (A-B) Dots plots showing Gene Set Enrichment Analysis (GSEA) of the royalblue module signature using (A) Gene Ontology (GO) and (B) Hallmark(1) gene sets where y-axis represents normalized enrichment scores (NES) in which scores greater than or less than zero represent gene sets enriched in genes positively or negatively correlated with the royalblue eigengene, respectively. The x-axis represents gene sets ordered by their enrichment score. Select top gene sets are labeled: Regulation of Wnt signaling (NES score 2.15, NES rank 14 of 6,033) in (A) and Epithelial Mesenchymal Transition (NES score 2.39, NES rank 1 of 50) in (B). Dots larger in size represent higher -log10(*P*) values. (C) Pink subnetwork where pink genes, royalblue genes, and known PAH genes (red nodes) from disease-gene databases (Comparative Toxicogenomics Database(1) and DisGeNET(1)) were projected onto the lung Bayesian regulatory network in Figure 3A. Larger size nodes represent hub genes where node size is proportional to -log10(FDR) as determined by Key Driver Analysis (1–3). Light grey nodes represent hub genes of the pink subnetwork that are not pink, royalblue, or red genes. Note this is the same subnetwork as in Figure 3B but with royalblue genes displayed here.

**Supplemental Figure 10:**
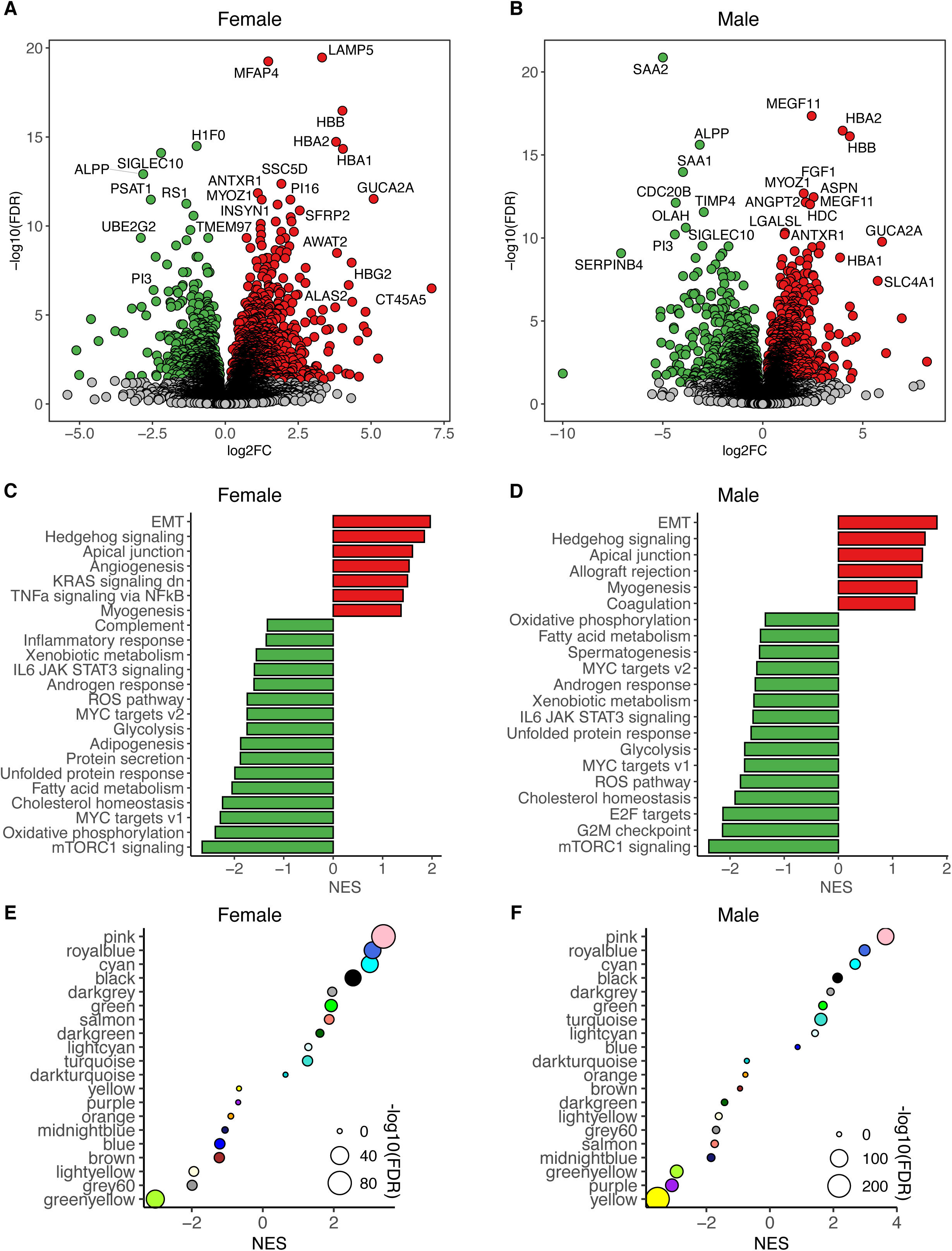
Sex-stratified analysis of dysregulated genes, pathways and modules. (A-B) Volcano plots showing upregulated genes in PAH lungs colored in red and downregulated genes colored in green among (A) females (71 PAH vs. 17 control) and (B) males (22 PAH vs. 34 control). Grey dots indicate genes with FDR ≥ 0.05. Select genes are labeled. (C-D) Bar plots showing GSEA results using the Hallmark pathway database and the DEG signature of PAH vs. control among (C) females and (D) males. Pathways enriched in genes upregulated in PAH with normalized enrichment score (NES) > 0 are colored in red and pathways enriched in downregulated genes with NES < 0 in green. Only pathways with FDR < 0.05 are shown. (E-F) Dot plots showing enrichment of modules as determined by GSEA for the PAH lung differential transcriptome among (E) females and (F) males. Larger size dots indicate stronger FDR value. log2FC = log2 fold change; NES = normalized enrichment score; EMT = epithelial-mesenchymal transition.

